# Eye opening differentially modulates inhibitory synaptic transmission in the developing visual cortex

**DOI:** 10.1101/199661

**Authors:** Wuqiang Guan, Jun-Wei Cao, Lin-Yun Liu, Zhi-Hao Zhao, Yinghui Fu, Yong-Chun Yu

**Affiliations:** Jing’an District Center Hospital of Shanghai, Institutes of Brain Science, State Key Laboratory of Medical Neurobiology and Collaborative Innovation Center for Brain Science, Fudan University, 131 Dong’An Road, Shanghai 200032, China

## Abstract

Eye opening, a natural and timed event during animal development, influences cortical circuit assembly and maturation; yet, little is known about its precise effect on inhibitory synaptic connections. Here we show that coinciding with eye opening, the strength of unitary inhibitory postsynaptic currents (uIPSCs) from somatostatin-expressing interneurons (SST-INs) to nearby excitatory neurons, but not interneurons, sharply decreases in layer 2/3 of the mouse visual cortex. In contrast, the strength of uIPSCs from fast-spiking interneurons (FS-INs) to excitatory neurons significantly increases during eye opening. More importantly, these developmental changes can be prevented by dark rearing or binocular lid suture, and reproduced by artificial opening of sutured lids. Mechanistically, this differential maturation of synaptic transmission is accompanied by a significant change in the postsynaptic quantal size. Together, our study reveals a differential regulation in GABAergic circuits in the cortex driven by eye opening likely crucial for cortical maturation and function.

## Introduction

During neocortical development, sensory experience critically influences neuronal connectivity and synaptic transmission (***Katz and Shatz, 1996***). In the visual system, maturation of the visual cortex depends on visual afferent activity (***Hensch, 2005***; ***Hofer et al., 2009***; ***Pecka et al., 2014***). The initial visual inputs experienced through closed eyelids are imprecise and diffuse, and followed by patterned visual inputs after eye opening. In general, experience-dependent maturation of the visual cortex is gradual, but the process accelerates soon after eye opening with the sudden and rapid increase in the intensity and frequency of afferent visual activity (***Lu and Constantine-Paton, 2004***). Eye opening in rodents typically occurs over a short period of one to two days (***Gandhi et al., 2005***). When experimentally synchronized, it drives a rapid series of changes in neuronal activity, protein trafficking, synaptogenesis, synaptic receptor composition, synaptic transmission and plasticity (***Lu and Constantine-Paton, 2004***; ***Yoshii et al., 2003***; ***Zhao et al., 2006***). Moreover, eye opening affects neuronal circuit development in the ascending retinothalamo-cortical pathway at every level: the retina (***Feller, 2003***; ***Tian and Copenhagen, 2003***), superior colliculus (SC) (***Zhao et al., 2006***), lateral geniculate nucleus (LGN) (***Hooks and Chen, 2006***; ***Levitt et al., 2001***), and visual cortex (***Hoy and Niell, 2015***; ***Ko et al., 2013, 2014***; ***Pecka et al., 2014***). For example, the probability and strength of excitatory connections between layer 2/3 pyramidal cells (PCs) in the visual cortex significantly increased after eye opening, and these changes were prevented by dark rearing (***Ishikawa et al., 2014***). Although previous study\ reported orientation tuning preferences of fast-spiking interneurons were dependent on normal visual experience after eye opening (***Kuhlman et al., 2011***), whether eye opening shapes inhibitory synaptic transmission in the neocortex during development remains unclear.

In the mature neocortex GABAergic interneurons can at a basic level be broken down into three main populations defined by expression of the calcium-binding protein parvalbumin, the neuropeptide somatostatin and the serotonin ionotropic receptor (***Lee et al., 2010***). Somatostatin-expressing interneurons (SST-INs), comprising approximately 20%∼30% of all neocortical interneurons, are one of the most prominent GABAergic interneuron subtypes in the neocortex (***Lee et al., 2010***; ***Pfeffer et al., 2013***). SST expression in the superficial neocortex has typically been associated with Martinotti cells (MCs)—GABAergic interneurons with ascending axons that arborize in cortical layer 1 and spread horizontally to neighboring columns (***Fino and Yuste, 2011***; ***Ma et al., 2006***). A large fraction of SST-INs preferentially target distal dendrites of PCs (***Cristo et al., 2004***). Cortical SST-INs provide lateral inhibition to local PC networks, and precisely control the efficacy and plasticity of glutamatergic inputs by regulating postsynaptic spine Ca^2+^ signals, synaptic dynamics, dendritic spike bursts, and transformation of dendritic inputs (***Chiu et al., 2013***; ***Higley, 2014***; ***Lovett-Barron et al., 2012***). Furthermore, cortical SST-INs not only innervate PCs, but also frequently inhibit other cortical interneurons (disinhibition) (***Pfeffer et al., 2013***). Parvalbumin-expressing (PV) fast-spiking (FS) INs, which constitute ∼40% of cortical GABAergic neurons, make powerful synapses onto the somatic and perisomatic compartments of PCs (***Cristo et al., 2004***). Although the emergence and maturation of connections from PV-INs to PCs and PV-INs have been reported previously ***(Lazarus and Huang, 2011***; ***Pangratz-Fuehrer and Hestrin, 2011***; ***Yang et al., 2014***), the development of inhibitory synaptic transmission (including amplitude and connectivity) from SST-INs to PCs and other interneuron subtypes remains largely unclear. In addition, accumulating evidence indicates that SST-INs regulate learning-induced and experience-dependent cortical plasticity through feedforward as well as feedback inhibitions, both during development and in adulthood (***Bloodgood et al., 2013***; ***Chen et al., 2015***; ***Marques-Smith et al., 2016***; ***Oh et al., 2016***; ***Tuncdemir et al., 2016***). There is also clear evidence to suggest that PV-INs regulate critical-period experience-dependent plasticity ***(Hensch, 2005***; ***Krishnan et al., 2015***). These observations raise important questions: Does eye opening play a role in regulation of inhibitory synaptic transmission from SST-INs and FS-INs to PCs or other types of interneurons, and if so, what is the synaptic mechanism underlying it?

In this study, we demonstrate that eye opening rapidly weakens the inhibitory synaptic transmission from SST-INs to PCs, whereas it increases inhibitory synaptic transmission from FS-INs to PCs. In addition, we show that the maturations of inhibitory synaptic transmission are mediated by differential changes in the postsynaptic quantal size.

## Results

### Rapid weakening of synaptic transmission from SST-INs onto PCs coincides with eye opening

To explicitly identify SST-INs in the neocortex, we crossed the *SST*-*IRES*-*Cre* mice with the loxP-flanked *Rosa26reporter*-*tdTomato* mice. The resulting progeny had SST-INs expressing red fluorescent tdTomato protein in the brain, which facilitated electrophysiological recordings (***Taniguchi et al., 2011***). We focused on layer 2/3 of the primary visual cortex. Consistent with previous reports (***Hu et al., 2013***; ***Pfeffer et al., 2013***), we observed ∼5.2% of tdTomato^+^ neurons expressed PV (5.2% ± 0.3%, 9 slices from 3 mice, ***Figure 1-figure supplement 1B***). Meanwhile, ∼17.2% of tdTomato^+^ neurons showed the fast-spiking properties (36/209, ***Figure 1-figure supplement 1C***) (***Hu et al., 2013***; ***Jiang et al., 2015***). Furthermore, FS tdTomato^+^ cells exhibited the distinctive basket cell morphology (4 out of 4, ***Figure 1-figure supplement 1C***). These cells were omitted from further analysis. Non-FS tdTomato^+^ cells were further characterized by morphological properties (***Figure 1-figure supplement 1A***). Typically, non-FS tdTomato^+^ neurons had ascending axonal arborizations with extensive branching in layer 1 and horizontal collaterals (19 out of 20, ***Figure 1-figure supplement 1A***). This morphology fits well with that of Martinotti cells, as previously described (***Fino and Yuste, 2011***).

To study synaptic transmission from SST-INs to PCs, we performed triple or quadruple whole-cell patch-clamp recordings to simultaneously record a layer 2/3 tdTomato^+^ SST-IN and two or three nearby PCs whose cell bodies were within ∼100 µm apart (***Figure 1A*** and ***1B***). The PCs were verified with morphological characteristics including pyramidal shape, large soma and thick primary dendrites decorated with spines, as well as firing properties (***Figure 1-figure supplement 3A***) (***Lazarus and Huang, 2011***; ***Schubert et al., 2001***). Once all recordings were established, serial action potentials (at least 20 trials) were triggered in tdTomato^+^ SST-IN and the inward unitary inhibitory postsynaptic currents (uIPSCs) were measured in PCs (***Figure 1C*** and ***1D***). To systematically study the development of synaptic transmission from SST-INs to PCs (SST-INs→PCs), we examined 391 SST-IN→PC pairs at different developmental stages. We found that SST-IN→PC connections emerged at postnatal day 7–8 (P7–8). The connection probability increased significantly from P7–8 to P9–11 (χ^2^ test, *p* = 0.0046; ***Figure 1E***) and remained largely comparable from P9 to P20 (χ^2^ test, *p* = 0.149; ***Figure 1E***). Interestingly, we observed a ∼65% reduction in the peak amplitude of uIPSCs from P12–13 to P14–15 (P7–8, 17.9 ± 2.9 pA; P9–11, 28.3 ± 5.4 pA; P12–13, 34.3 ± 4.8 pA; P14–15, 11.6 ± 1.8 pA; P16–17, 10.0 ± 1.4 pA; P18–20, 6.0 ± 1.9 pA; one-way ANOVA, *F*(5,185) *=* 8.788, *p* = 1.7 × 10^-7^; ***Figure 1F***). Notably, this dramatic change coincided with the natural eye openings of mice under standard living conditions (***Figure 1*-*figure supplement 2***), suggesting that the weakening of SST-IN→PC uIPSCs may result from eye opening. Although the 10%-90% rise time of uIPSCs at P18–20 was significantly larger than that at P7–8 and P12–13, the 10%-90% rise time of uIPSCs did not significantly change at P9–17 (***Figure 1G***). Furthermore, the half-width of uIPSCs did not show any obvious change at P7–20 (***Figure 1H***). In addition, the total length and complexity of both apical and basal dendrites of PCs exhibited no significant difference between P12–13 and P14–15 mice (***Figure 1-figure supplement 3B***n***3I***), suggesting that dendritic morphology of PCs are not changed during eye opening. Of note, the connection probability and strength of SST-IN→PC synaptic transmission exhibited similar development properties when we used cesium-based and high Cl^-^ internal solution to record the postsynaptic currents (***Figure 1*-*figure supplement 4***).

**FIGURE 1:**
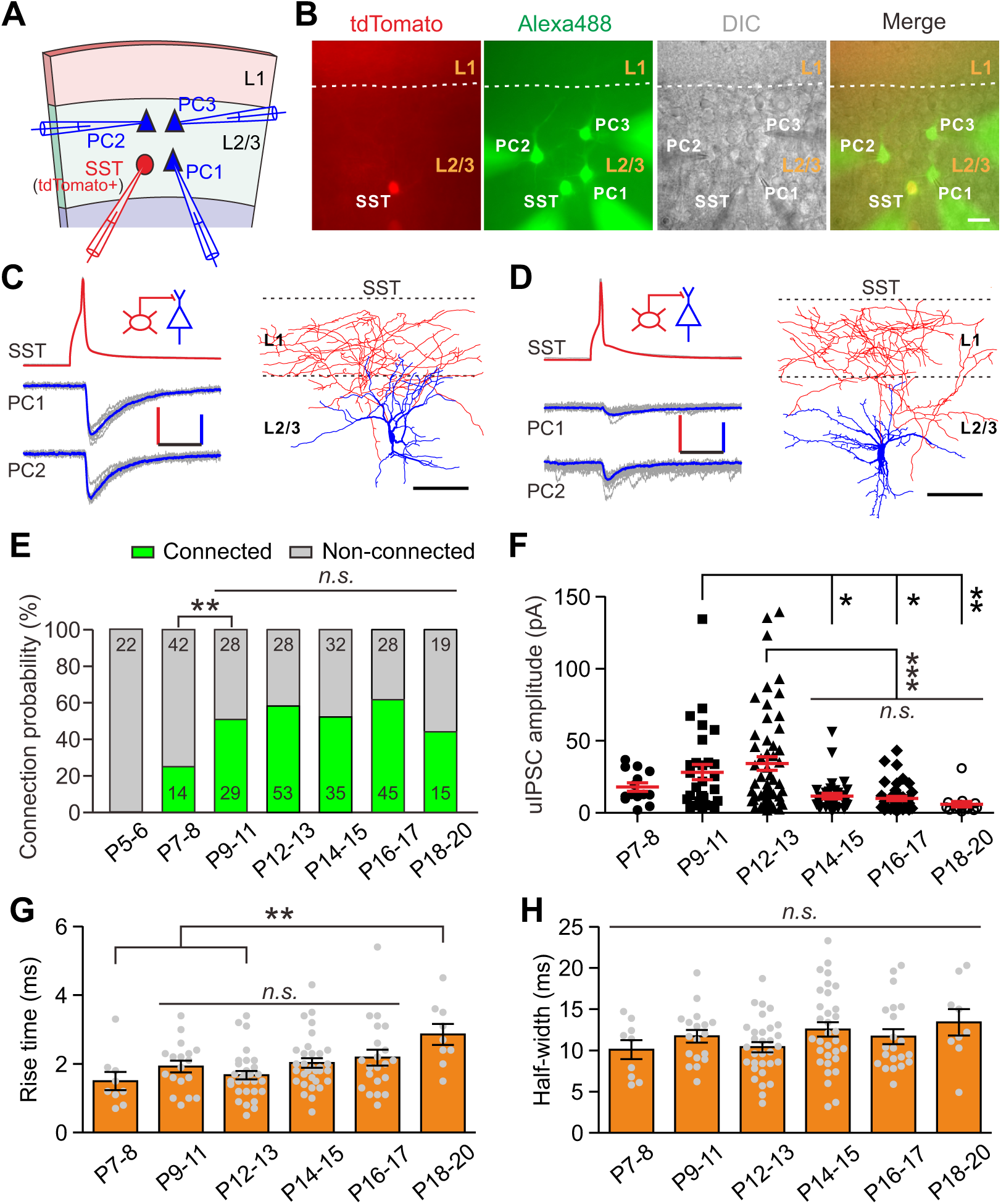
Development of synaptic transmission from SST-INs onto PCs in layer 2/3 of the visual cortex. (**A**) Schema of a quadruple whole-cell recording from a layer 2/3 SST-IN and three layer 2/3 (**B**) Representative fluorescent (tdTomato, SST-INs; Alexa 488, recording neurons), and merged images of a quadruple recording of a SST-IN and three PCs. The broken indicate the border of layer 1 and layer 2/3. Scale bar, 20 μm. (**C**) Left, representative showing synaptic transmission from a layer 2/3 SST-IN to two layer 2/3 PCs recorded The red and blue lines indicate the averaged traces. Scale bars: 50 mV (vertical, red), (vertical, blue) and 20 ms (horizontal). Right, morphological reconstruction of the Scale bar: 80 µm. (D) Left, representative traces showing synaptic transmission a layer 2/3 SST-IN to two layer 2/3 PCs recorded at P15. Scale bars: 50 mV (vertical, 25 pA (vertical, blue) and 20 ms (horizontal). Right, morphological reconstruction of SST-IN. Scale bar: 80 µm. (**E**) Probability of synaptic connection from SST-INs to PCs P5–20. Numbers in the bar indicate the number of pairs in each connection category. In 391 pairs were recorded from 82 mice. (**F**) Quantification of the peak amplitude of from SST-INs to PCs at different postnatal ages. (**G**-**H**) Quantification of the 10%-rise time (**G**) and half-width (**H**) of uIPSCs at P7–20. Detailed statistical analysis, data and exact sample numbers are presented in ***Figure 1*-*source data 1.*** Error bars indicate mean ± SEM. *, *p* < 0.05; **, *p* < 0.01; ***, *p* < 0.001; *n.s.* for *p* > 0.05.

To assess whether other cortical areas undergo similar developmental changes, the connection probability and strength of uIPSCs from SST-INs to PCs were measured in layer 2/3 of the prefrontal cingulate cortex area 1/2 (Cg1/2) during P9–20 (***Figure 2***). The connection probability remained comparable from P9 to P20 (χ^2^ test, *p* = 0.857; ***Figure 2B***). Consistent with the visual cortex, the peak amplitude of SST-IN→PC uIPSCs at P14–15 was significantly smaller than that at P12–13 in Cg1/2 (P9–11, 31.6 ± 6.6 pA; P12–13, 31.0 ± 4.7 pA; P14–15, 11.0 ± 3.1 pA; P16–17, 5.0 ± 1.9 pA; P18n 20, 5.7 ± 1.0 pA; one-way ANOVA, *F*_(4,46)_ *=* 5.51, *p* = 6.0 × 10^-4^; ***Figure 2C***). These results suggest that the weakening of SST-IN→PC synaptic transmission during eye opening exists not only in the visual cortex, but also in Cg1/2.

**FIGURE 2:**
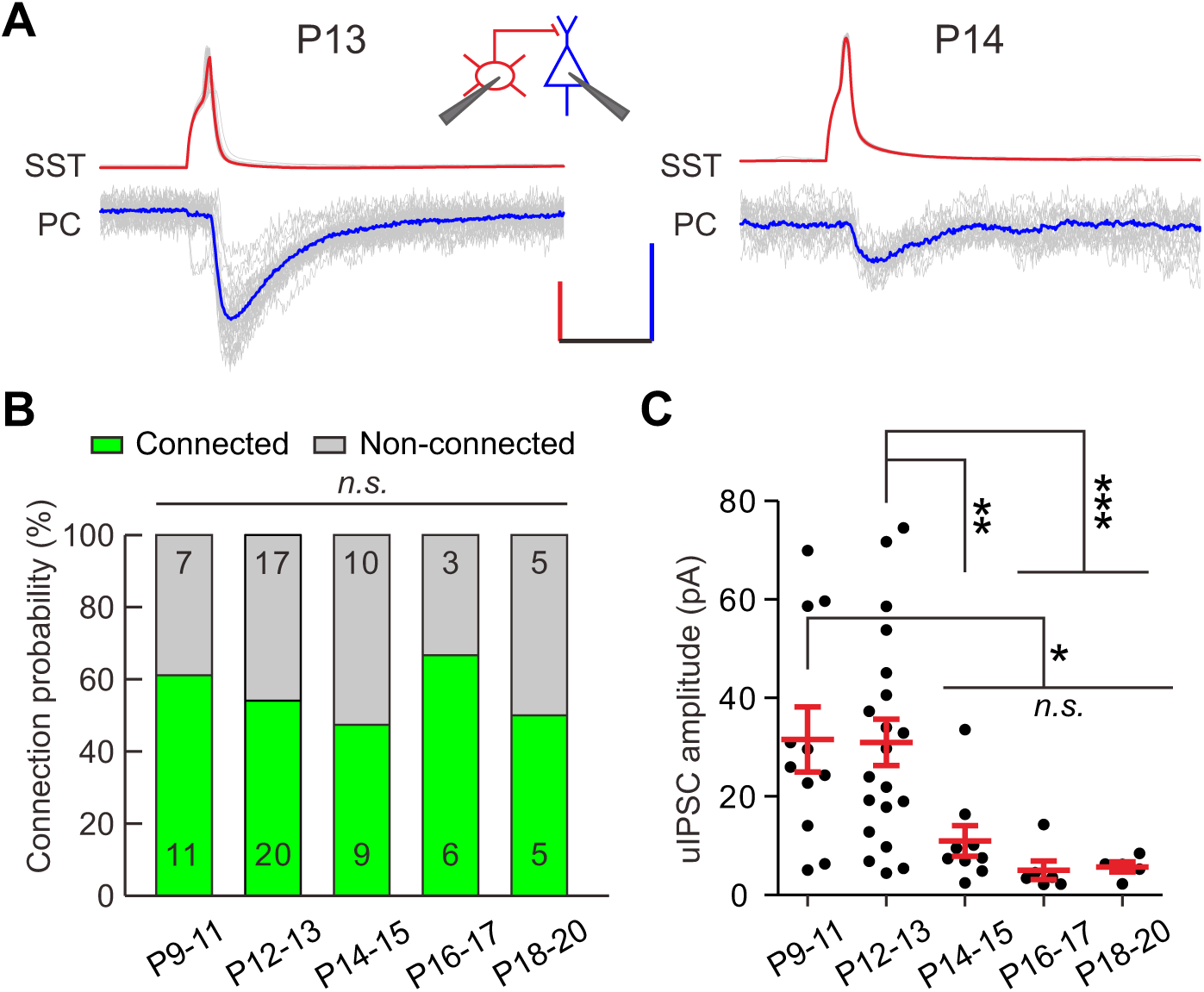
>Development of synaptic transmission from SST-INs onto PCs in prefrontal Cg1/2 area. (**A**) Representative traces of synaptic transmission from a layer 2/3 SST-IN to a layer 2/3 PC prefrontal Cg1/2 area at P13 and P14. Inset schema indicates paired patch recording of a (red) and a PC (blue). Scale bars: 50 mV (vertical, red), 20 pA (vertical, blue) and ms (horizontal). (**B**) Histogram of the connection probability from P9 to P20. (**C**) of the peak amplitude of uIPSCs from P9 to P20. Detailed statistical analysis, data and number of experiments are presented in the ***Figure 2*-*source data 1***. *, *p* < *p* < 0.01; ***, *p* < 0.001; *n.s.* for *p* > 0.05.

Together, these results demonstrate that coinciding with eye opening, SST-IN→PC synaptic transmission dramatically weakens in cortical layer 2/3.

### FS-IN→PC synaptic transmission increases during eye opening

We further examined whether the strength of synaptic transmission from fast-spiking PV interneurons (FS-INs) to PCs (FS-INs→PCs) could change during eye opening. We took advantage of *Lhx6*-*EGFP* transgenic mice, in which the majority of MGE-derived interneurons are labeled, and bred this line onto *SST*-*tdTomato* line (*SST*-*tdTomato::Lhx6*-*EGFP* line), allowing us to distinguish between SST-INs (tdTomato^+^) and other types of interneurons (EGFP^+^/tdTomato^-^) (***Figure 3A***) (***Tuncdemir et al., 2016***). EGFP^+^/tdTomato^-^ FS-INs were further determined with the fast-spiking properties (***Figure 3C*** and ***Figure 3-figure supplement 1***). We found that 78.4% of recorded EGFP^+^/tdTomato^-^ cells were FS-INs at P12–18 (87 out of 111, ***Figure 3-figure supplement 1***), and these FS-INs showed basket cell morphology (15 out of 15, ***Figure 3-figure supplement 1***). To examine FS-IN→PC synaptic transmission, a layer 2/3 EGFP^+^/tdTomato^-^ FS-IN and nearby tdTomato^-^/EGFP^-^ PCs were simultaneously recorded (***Figure 3B*** and ***3C***). The connection probability did not significantly change from P12 to P18 (χ^2^ test, *p* = 0.949; ***Figure 3D***). Interestingly, unlike SST-IN→PC synaptic transmission, the strength of FS-IN→PC uIPSCs at P14–15 and P16–18 was significantly larger than that at P12–13 (P12–13, 91 ± 14 pA; P14–15, 217 ± 37 pA; P16–18, 279 ± 57 pA; one-way ANOVA, *F*_(2,63)_ = 8.71, *p* = 4.6 × 10^-4^; ***Figure 3E***), suggesting that eye opening may increase the FS-IN→PC synaptic transmission. Moreover, the 10%n90% rise time and half-width of uIPSCs did not exhibit obvious change between P12–13 and P14–15, while the rise time was significantly smaller at P16–18 than at P12–13 (***Figure 3F*** and ***3G***). Together, these results demonstrate that FS-IN→PC synaptic transmission increases during eye opening.

**FIGURE 3:**
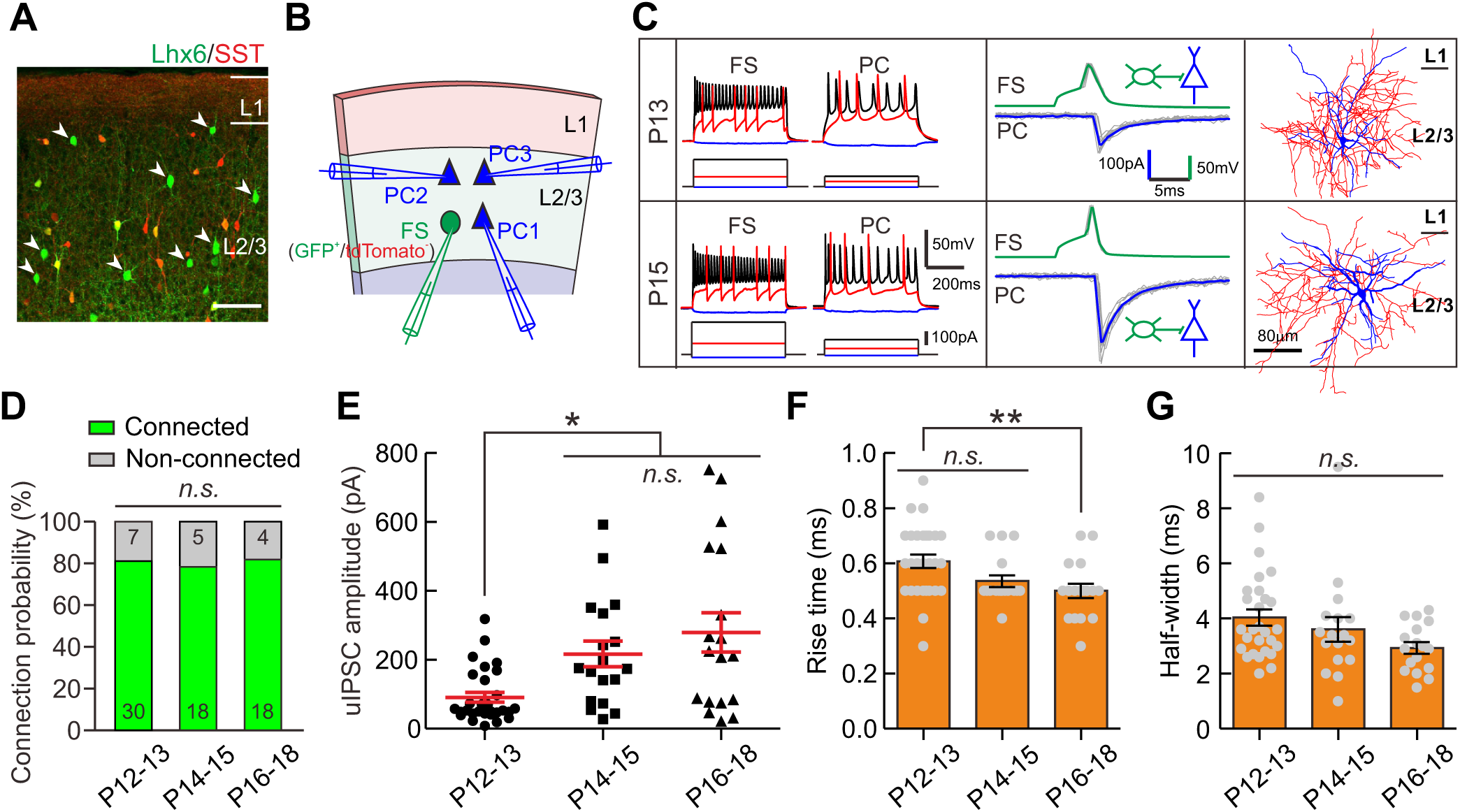
Strength of synaptic transmission from FS-INs onto PCs significantly increases in layer 2/3 of the visual cortex during eye opening. (**A**) Fluorescent image of a visual coronal section from *SST*-*tdTomato::Lhx6*-*EGFP* line. + - SST-INs; GFP, Lhx6-EGFP cells. Arrow heads indicate EGFP /tdTomato cells. bar, 100 μm. (**B**) Schema of a quadruple whole-cell recording from a layer 2/3 FS-IN + - /tdTomato) and three layer 2/3 PCs. (**C**) Two examples of connection from a layer FS-IN to a layer 2/3 PC at P13 and P15. Left panels, membrane potential responses of FS-INs and PCs to current injections. Middle panels, synaptic transmission from to PCs. Right panels, reconstructed morphology of recorded FS-INs. (**D**) The connection probability from FS-INs to PCs did not change from P12 to P18. (**E**) Peak of FS-IN PC uIPSCs at P14–15 and P16–18 was significantly larger than that at (**F**-**G**) Quantification of the 10%-90% rise time (**F**) and half-width (**G**) of uIPSCs at 18. Detailed statistical analysis, detailed data and number of experiments are presented in the ***Figure 3*-*source data 1***. *, *p* < 0.05; **, *p* < 0.01; *n.s.* for *p* > 0.05.

### Synaptic transmission from SST-INs to other types of interneurons does not change during eye opening

SST-INs not only frequently innervate pyramidal neurons, but also strongly inhibit other types of interneurons (***Pfeffer et al., 2013***). We first examined the development of GABAergic synaptic transmission from SST-INs to FS-INs (SST-INs→FS-INs) in *SST*-*tdTomato::Lhx6*-*EGFP* line during eye opening. To study SST-IN→FS-IN synaptic transmission, we simultaneously recorded from a layer 2/3 tdTomato^+^ SST-IN and a nearby EGFP^+^/tdTomato^-^ FS-IN (***Figure 4A***). We compared the connection probability and strength of SST-IN→FS-IN uIPSCs at P12–13, P14–15 and P16–18 (***Figure 4B*** and ***4C***). Our data showed that the SST-IN→FS-IN connection probability did not change from P12 to P18 (χ^2^ test, *p* = 0.855; ***Figure 4B***). Notably, the strength of SST-IN→FS-IN uIPSCs did not exhibit any obvious change (P12–13, 62.0 ± 12.7 pA; P14–15, 72.1 ± 18.7 pA; P16–18, 56.6 ± 13.0 pA; one-way ANOVA, *F*_(2,24)_ = 0.268, *p* = 0.767, ***Figure 4C***). These results suggest that the strength of SST-IN→FS-IN synaptic transmission does not change during eye opening.

**FIGURE 4:**
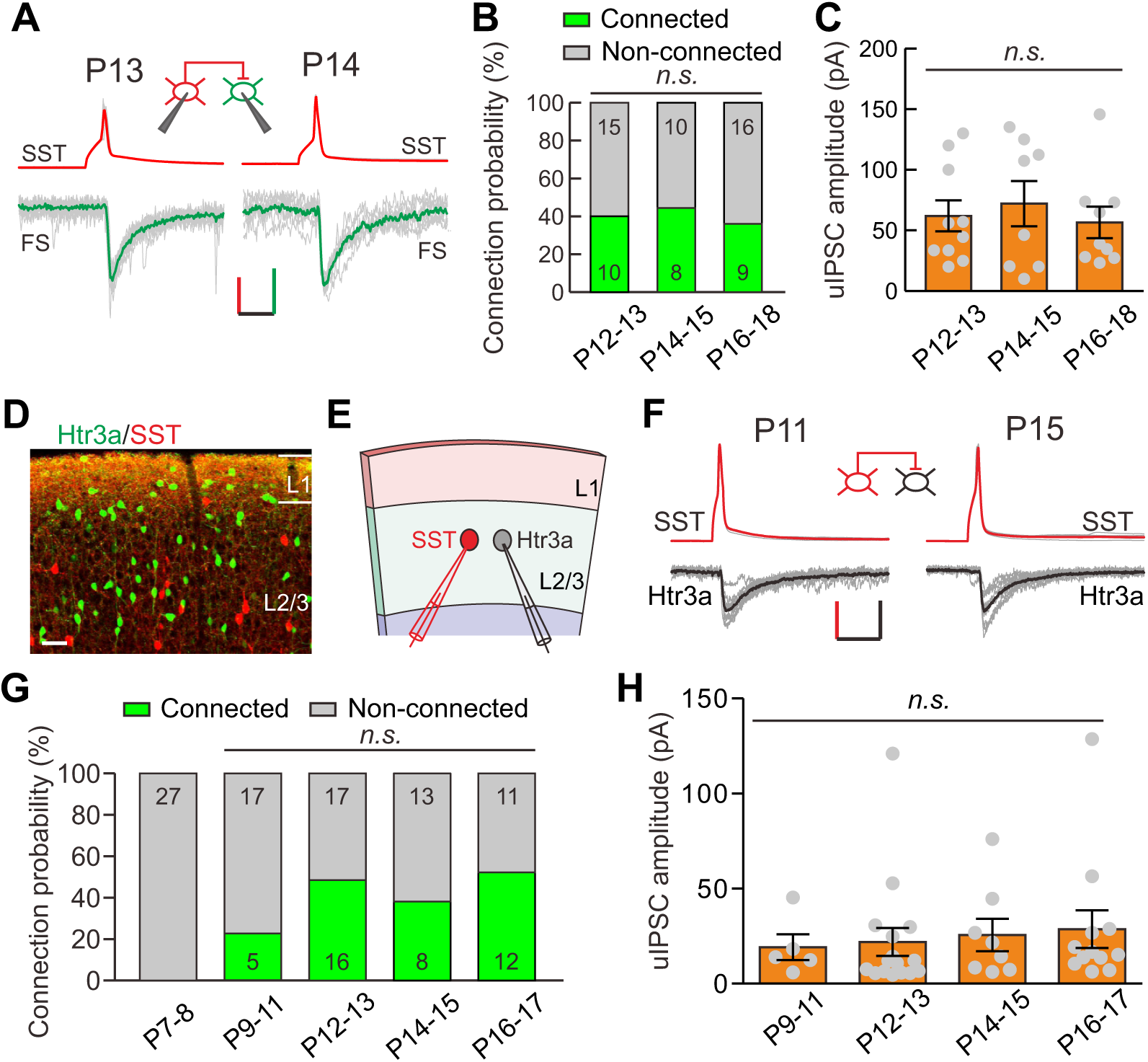
Development of synaptic transmission from SST-INs to FS-INs and Htr3a-INs. (**A**) Representative evoked responses from a layer 2/3 SST-IN to a FS-IN at P13 and P14, respectively. Inset panel: schematic of a paired recording from a SST-IN (red) and a FS-IN (green). bars: 50 mV (vertical, red), 25 pA (vertical, green) and 20 ms (horizontal). (**B**) Summary of probability from layer 2/3 SST-INs to FS-INs at different postnatal ages. In total, 68 were recorded from 11 mice. (**C**) The peak amplitude of uIPSCs from layer 2/3 SST-INs to was unchanged from P12 to P18. (**D**) Fluorescent image of a visual coronal section from *tdTomato::Htr3a*-*EGFP* line. TdTomato, SST-INs; GFP, Htr3a-INs. Scale bar, 50 μm. (**E**) of a paired recording from a layer 2/3 SST-IN and a layer 2/3 Htr3a-IN. (**F**) Traces representative synaptic transmission from a layer 2/3 SST-IN to a layer 2/3 Htr3a-IN at and P15. Scale bars: 40 mV (vertical, red), 20 pA (vertical, black) and 20 ms (horizontal). (**G**) of connection probability from layer 2/3 SST-INs to layer 2/3 Htr3a-INs at different ages. In total, 126 pairs were recorded from 17 mice. (**H**) The peak amplitude of uIPSCs layer 2/3 SST-INs to Htr3a-INs did not significantly change from P9 to P17. Detailed statistical analysis, detailed data and exact sample numbers are presented in ***Figure 4*-*source data 1***. Error bars indicate mean ± SEM. *n.s.* for *p* > 0.05.

We next explored the development of synaptic transmission from layer 2/3 SST-INs to layer 2/3 Htr3a-positive interneurons (Htr3a-INs). Htr3a-INs are the predominant type of interneurons in the visual superficial layers, comprising ∼50% of layer 2/3 interneurons (***Lee et al., 2010***). To simultaneously label SST-INs and Htr3a-INs, we crossed heterozygous *SST*-*tdTomato* mice with *Htr3a*-*EGFP* mice (*SST*-*tdTomato::Htr3a*-*EGFP* line). Previous studies have demonstrated that 5HT3a receptors (encoded by *Htr3a*) in the neocortex are present exclusively in GABAergic interneurons (***Lee et al., 2010***). In *SST*-*tdTomato::Htr3a*-*EGFP* mice, SST-INs expressed tdTomato, while Htr3a-INs expressed EGFP (***Figure 4D***). Only ∼1% of tdTomato^+^ SST-INs expressed EGFP (1.1% ± 0.7%, 6 slices from 3 mice), suggesting that SST-INs and Htr3a-INs belong to two different types of interneurons, as previously reported (***Chittajallu et al., 2013***; ***Lee et al., 2010***). To assess SST-IN→Htr3a-IN synaptic transmission, simultaneous recordings were made on pairs of tdTomato^+^ SST-INs and GFP^+^ Htr3a-INs in layer 2/3 of the primary visual cortex (***Figure 4E*** and ***4F***). SST-IN→Htr3a-IN connections emerged at P9–11, and the probability did not change significantly from P9 to P17 (χ^2^ test, *p* = 0.169; ***Figure 4G***). Similar to SST-IN→FS-IN uIPSCs, there was no obvious difference in the strength of SST-IN→Htr3a-IN uIPSCs between P12–13 (before eye opening) and P14–15 (after eye opening) mice (P9–11, 19.1 ± 6.8 pA; P12–13, 21.9 ± 7.4 pA; P14–15, 25.6 ± 8.5 pA; P16–17, 28.6 ± 9.9 pA; one-way ANOVA, *F*_(3,37)_ = 0.183, *p* = 0.907; ***Figure 4H***). These results suggest that the strength of SST-IN→Htr3a-IN synaptic transmission does not change during eye opening.

Taken together, these results show that the strength of synaptic transmission from SST-INs to PCs, but not from SST-INs to other types of interneurons, selectively decreases in layer 2/3 of the cortex upon eye opening. To further confirm the time and target cell type differential inhibition by SST-INs, we performed triple recordings to simultaneously record from a layer 2/3 SST-IN and both a PC and a FS-IN in *SST*-*tdTomato:: Lhx6*-*EGFP* mice (***Figure 5A***), or from a layer 2/3 SST-IN and both a PC and a layer 2/3 Htr3a-IN in *SST*-*tdTomato::Htr3a*-*EGFP* mice (***Figure 5F***), at P12–13 and P14–15. The uIPSCs recorded simultaneously from two different postsynaptic cell types were directly compared. SST-IN inhibition onto PCs and FS-INs exhibited no significant difference at P12–13 (paired *t* test, *p* = 0.510; ***Figure 5B, 5C*** and ***5E***). However, at P14–15, the same SST-INs produced much larger uIPSCs in FS-INs than in PCs (paired *t* test, *p* = 0.037; ***Figure 5B, 5D*** and ***5E***). Similarly, SST-IN inhibition onto PCs and layer 2/3 Htr3a-INs was not significantly different at P12–13 (paired *t* test, *p* = 0.601; ***Figure 5G*** and ***5I***), but SST-INs produced significantly larger uIPSCs in layer 2/3 Htr3a-INs than in PCs at P14–15 (paired *t* test, *p* = 0.037; ***Figure 5H*** and ***5I***). These results suggest that the inhibitory synapses made by SST-INs exhibit remarkable specificity in their connection strength with specific targets before and after eye opening.

**FIGURE 5:**
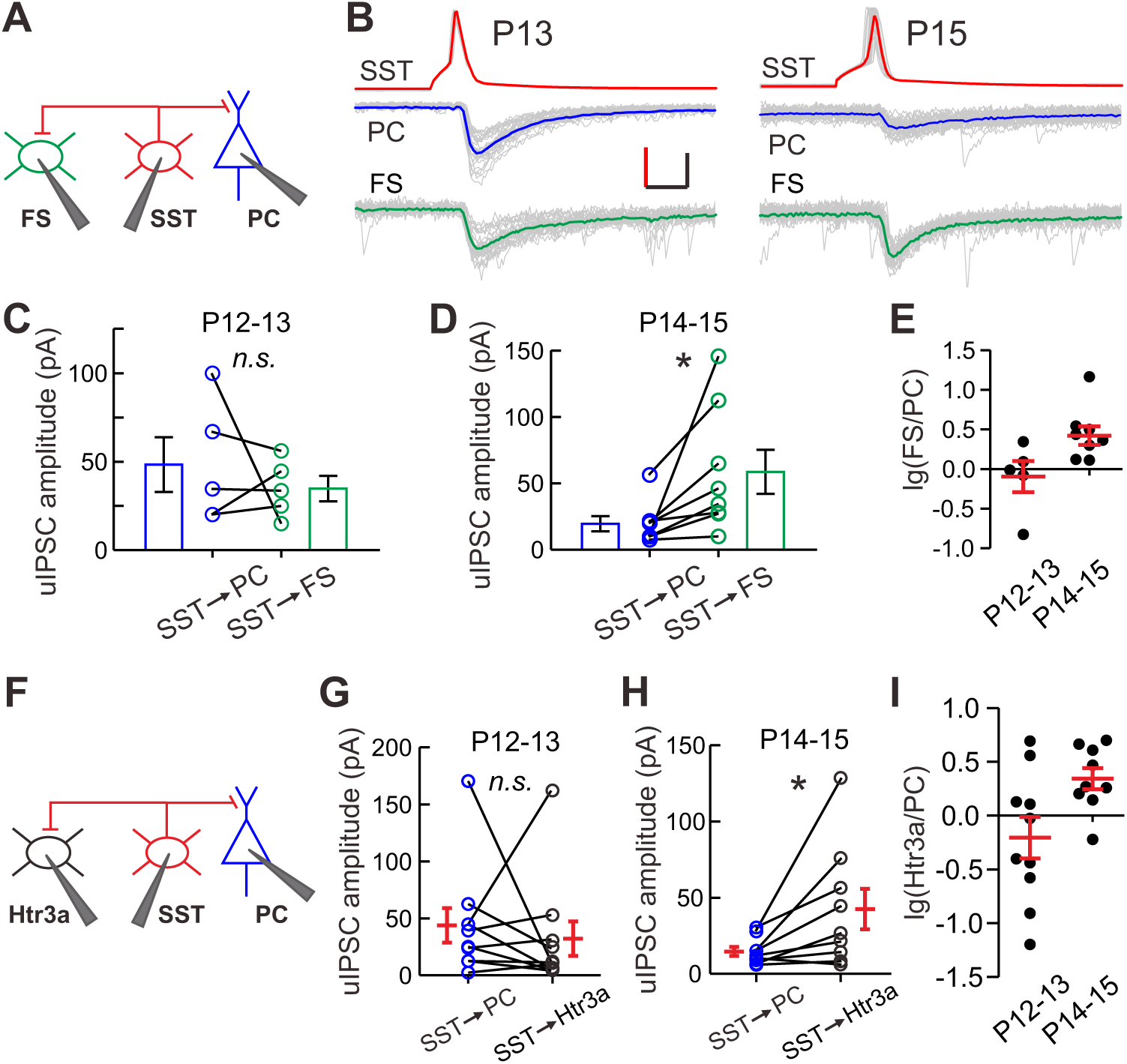
Target cell type differential inhibition by SST-INs before and after eye opening. (**A**) Schematic of a triple recording from a layer 2/3 SST-IN, a PC and a layer 2/3 FS-IN. (**B**) Representative evoked responses from a layer 2/3 SST-IN simultaneously onto a layer 2/3 PC and a 2/3 FS-IN at P13 and P15. Scale bars: 50 mV (vertical, red), 50 pA (vertical, black) and 5 ms (horizontal). (**C**) uIPSC amplitude evoked by SST-INs was not significantly different between layer PCs and FS-INs at P12–13. (**D**) uIPSC amplitude evoked by SST-INs was significantly smaller than in FS-INs at P14–15. (**E**) Logarithm of the ratio between uIPSC amplitude in FS-INs and PCs at P12–13 and P14–15. (**F**) Schematic of a triple recording from a layer 2/3 SST-IN, a layer PC and a layer 2/3 Htr3a-IN. (**G**) uIPSC amplitude evoked by SST-INs was not significantly between PCs and Htr3a-INs at P12–13. (**H**) uIPSC amplitude evoked by SST-INs was significantly smaller in PCs than in Htr3a-INs at P14–15. (**I**) Logarithm of the ratio between uIPSC in Htr3a-INs and PCs at P12–13 and P14–15. Detailed statistical analysis, detailed data exact sample numbers are presented in the ***Figure 5*-*source data 1***. Error bars indicate mean ± *, *p* < 0.05; *n.s.* for *p* > 0.05.

### Eye opening modulates the SST-IN→PC and FS-IN→PC synaptic transmission

Although the weakening of synaptic transmission from SST-INs to PCs was observed at the time of eye opening, it could be induced by intrinsic developmental programs or other developmental programs with coincidental timing rather than due to eye opening per se. To determine which factor is responsible, we first deprived the visual inputs by dark rearing. We dark-reared *SST*-*tdTomato* mice from P3, and recorded the synaptic transmission from SST-INs to PCs in layer 2/3 of the primary visual cortex at P12–15 (***Figure 6-figure supplement 1A***). Under dark rearing condition, no significant changes were observed in both connection probability (χ^2^ test, *p* = 0.066; ***Figure 6-figure supplement 1C***) or unitary strength of SST-IN→PC synaptic transmission between P12–13 and P14–15 (P12–13, 19.6 ± 4.1 pA, n = 13; P14–15, 21.7 ± 3.6 pA, n = 16; two-tailed unpaired *t* test, *p* = 0.702; ***Figure 6-figure supplement 1B*** and ***1D***). These results suggest that visual deprivation prevents the weakening of SST-IN→PC synaptic transmission at the time of eye opening.

Visual deprivation achieved by dark rearing not only blocks eye opening-induced visual inputs, but also eliminates the natural diffuse dark/light stimulation presented through the eyelids before eye opening. Therefore, it is difficult to determine whether elimination of the weakening of SST-IN→PC synaptic transmission is induced by eye opening deprivation, or by visual deprivation. To address this, we performed binocular lid suture from P8 to block eye opening, and manipulated the time of eye opening by artificially opening the lids in the binocular lid-sutured mice at P16 (two days after natural eye opening) (***Figure 6A***). The connection probability and strength of SST-IN→PC and FS-IN→PC synaptic transmission were further compared in the sutured mice at P12–13, P14–15 and P17–20, as well as artificially opened mice at P17–20 (i.e., different groups) (***Figure 6*** and ***Figure 6-figure supplement 2***).

**FIGURE 6:**
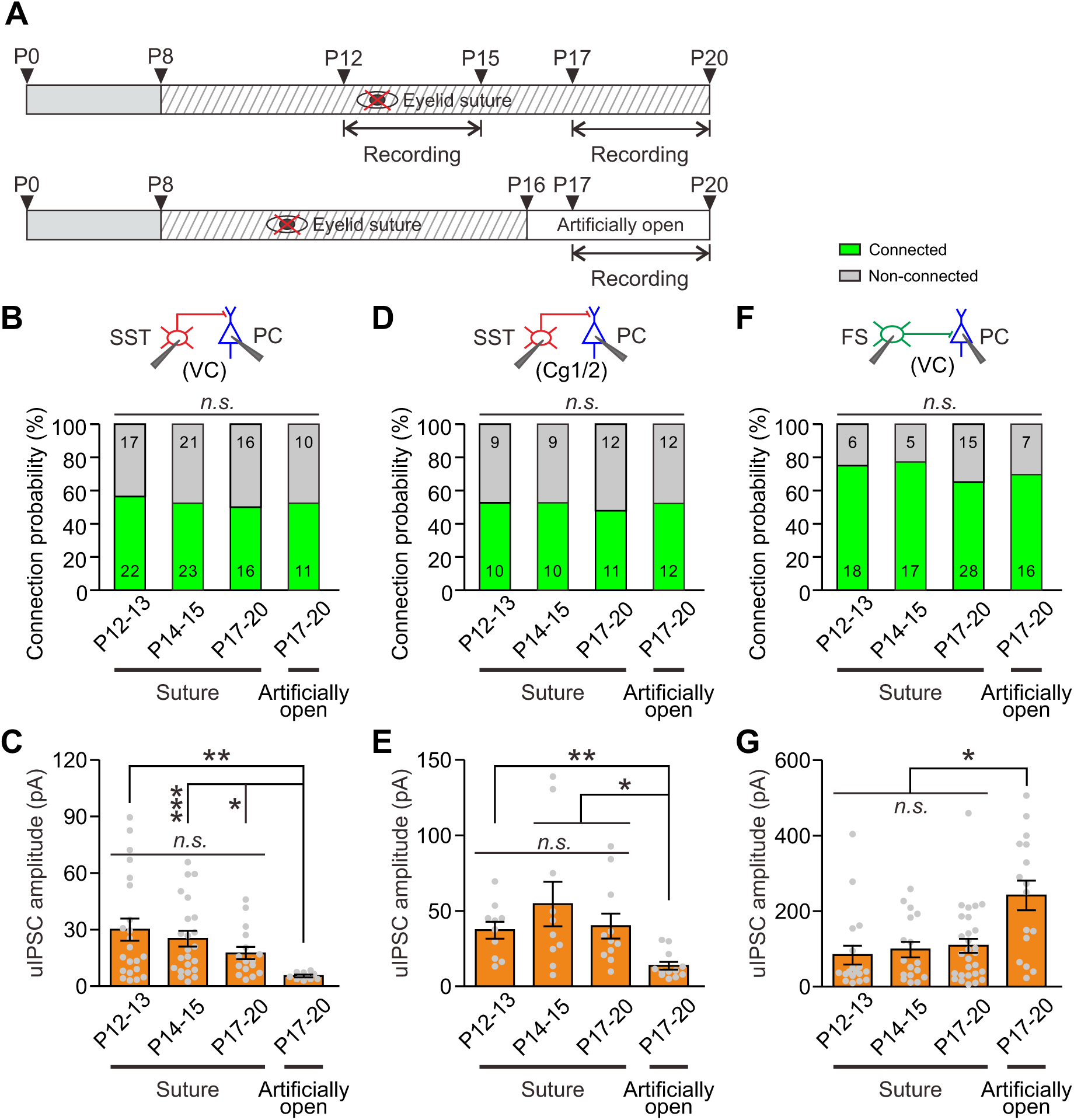
Eye opening modulates the strength of synaptic transmission from SST-INs to PCs and from FS-INs to PCs. (**A**)Schematic time schedule of eyelid suturing, eyelid reopening, and recording of synaptic (**B**) Summary of SST→IN PC connection probability in visual cortex (VC). (**C**) Quantification of the peak amplitude of SST→IN PC uIPSCs in VC. (**D**) Summary of SST→IN PC probability in Cg1/2. (**E**) Quantification of the peak amplitude of SST→IN PC uIPSCs Cg1/2. (**F**) Summary of FS→IN PC connection probability in VC. (**G**) Quantification of the peak of FS→IN PC uIPSCs in VC. Detailed statistical analysis, detailed data and exact sample are presented in the ***Figure 6*-*source data 1.*** Error bars indicate mean ± SEM. *, *p* < 0.05; < 0.01; *n.s.* for *p* > 0.05.

For SST-IN→PC synaptic transmission within the visual cortex (VC), the connection probability was not significantly different among the different groups (χ^2^ test, *p* = 0.958; ***Figure 6B***). Consistent with dark rearing, there was no significant change in the peak amplitude of uIPSCs between P12–13 and P14–15 sutured mice (***Figure 6C***). Moreover, the strength of SST-IN→PC uIPSCs at P17–20 in continuously sutured mice was comparable to those at P12–13 and P14–15 in sutured mice (sutured, P12–13, 30.1 ± 5.9 pA; sutured, P14–15, 25.2 ± 4.1 pA; sutured, P17–20, 17.5 ± 10.6 pA; ***Figure 6C***). These results suggest that eye opening deprivation prevents the weakening of SST-IN→PC synaptic transmission. Interestingly, the strength of SST-IN→PC uIPSCs in artificially opened mice at P17–20 was significantly lower than that in sutured mice at P12–13, P14–15 and P17–20 (one-way ANOVA, *F*(3,68) = 4.231, *p* = 0.008; ***Figure 6C***). These results suggest that artificial eye opening can induce the weakening of SST-IN→PC synaptic transmission and that this change associated with eye opening is unlikely due to an intrinsic developmental program. Notably, similar results were obtained in the prefrontal Cg1/2 area. In Cg1/2, the probability of SST-IN→PC connections was not significantly changed among different groups (χ^2^ test, *p* = 0.987; ***Figure 6D***). The strength of SST-IN→PC uIPSCs remained unchanged at sutured mice at P12–13, P14–15 and P17–20 (***Figure 6E***). Furthermore, the strength of SST-IN→PC uIPSCs in artificially eye opened mice at P17–20 were significantly smaller than that in sutured mice at P12–13, P14–15 and P17–20 (sutured: P12–13, 37.3 ± 5.7 pA, P14–15 54.6 ± 14.7 pA, P17–20, 39.9 ± 8.3 pA; artificially open: P17–20, 13.6 ± 2.6 pA; one-way ANOVA, *F*_(3,39)_ = 5.607, *p* = 0.003; ***Figure 6E***). In addition, both the connection probability and strength of SST-IN→FS-IN synaptic transmission were comparable at P12–13 and P14–15 in sutured mice (***Figure 6-figure supplement 3***).

For FS-IN→PC synaptic transmission in VC, similar to SST-IN→PC synaptic transmission, there were no significant changes in the connection probability among the different groups (χ^2^ test, *p* = 0.722; ***Figure 6F***). Likewise, the strength of FS-IN→PC uIPSCs was similar in the sutured mice at P12–13, P14–15 and P17–20; whereas the strength in artificially opened mice at P17–20 was significantly larger than that in sutured mice at P12–13, P14–15 and P17–20 (sutured: P12–13, 84 ± 25 pA, P14–15, 98 ± 21 pA, P17–20, 108 ± 19 pA; artificially open, P17–20, 241 ± 39 pA; one-way ANAVA, *F*(3,75) = 7.096, *p* = 2.9 × 10^-4^; ***Figure 6G***).

Together, these results strongly suggest that eye opening differentially modulates the strength of SST-IN→PC and FS-IN→PC synaptic transmission.

### Eye opening alters postsynaptic quantal size

We next examined presynaptic or postsynaptic mechanisms that underlie the differential changes of SST-IN→PC and FS-IN→PC synaptic transmission during eye opening. We assessed the presynaptic release probability by analysis of paired-pulse ratio (PPR), coefficient of variation (C.V.) and failure rate (***Miao et al., 2016***; ***Pouzat and Hestrin, 1997***). In SST-IN→PC synaptic transmission, there were no significant differences in PPR (two-way ANOVA, *F*(2,132) = 0.172, *p* = 0.842; ***Figure 7A*** and ***7B***), C.V. (P12–13, 0.494 ± 0.052, n = 20; P14–15, 0.472 ± 0.052, n = 23; two-tailedunpaired *t* test, *p* = 0.775; ***Figure 7C***) or failure rate (P12–13, 4.8% ± 2.2%, n = 20; P14–15, 5.3% ± 2.8%, n = 23; two-tailed unpaired *t* test, *p* = 0.897; ***Figure 7D***) before and after eye opening at P12–13 and at P14–15. These results suggest that eye opening does not affect the presynaptic release probability in SST-IN→PC synaptic transmission. The number of the presynaptic release sites (N) and the postsynaptic quantal size (Q) were estimated by the variance-mean (V-M) analysis (***Mitra et al., 2012***; ***Scheuss and Neher, 2001***). In brief, we used the theoretically expected parabolic relationship between the variance and mean of synaptic responses under multiple-pulse stimulation and different external Ca^2+^/Mg^2+^ concentrations (1 mM Ca^2+^/3 mM Mg^2+^, 2 mM Ca^2+^/2 mM Mg^2+^, 3.7 mM Ca^2+^/0.3 mM Mg^2+^) to obtain estimates of N and Q (***Mitra et al., 2012***). A train of 3 action potentials (20 Hz, 30–40 trials) was elicited in presynaptic SST-IN, and postsynaptic responses were recorded in PCs (***Figure 7E***). We used a Cs-based intracellular solution containing high concentration of Cl- (60 mM) to record the postsynaptic currents. The relationship between mean and variance under different external Ca^2+^/Mg^2+^ concentrations was plotted with the parabola fitness (see Materials and methods, ***Figure 7F***). We found that N in presynaptic terminals at P12–13 was similar to that at P14–15 (P12–13, 12.3 ± 1.5, n = 12; P14–15, 14.3 ± 1.9, n = 8; two-tailed unpaired *t* test, *p* = 0.419, figure ***7G***). However, Q at P14–15 was significantly lower than that at P12–13 (P12–13, 20.2 ± 2.2 pA, n = 12; P14–15, 8.7 ± 0.9 pA, n = 8; Mann Whitney *U* test, *p* = 2.9 × 10^-4^; ***Figure 7H***), suggesting that postsynaptic mechanisms contribute to the developmental weakening of SST-IN→PC synaptic strength.

**FIGURE 7:**
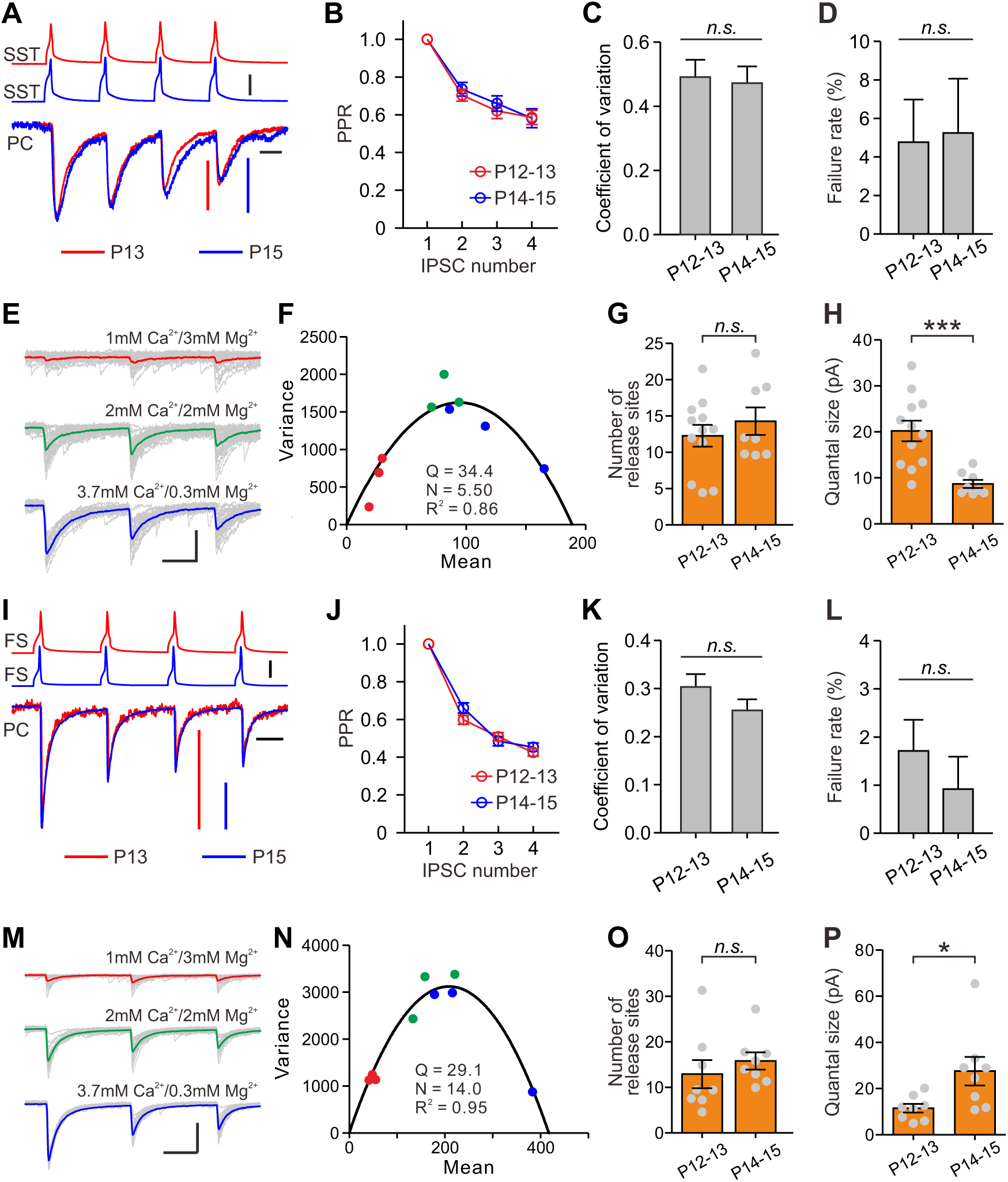
Postsynaptic mechanisms underlying the changes of synaptic transmission from SST-INs to PCs and from FS-INs to. (**A**) Amplitude-scaled overlay of paired-pulse ratio (PPR) responses in SST-IN→PC connections at P13 and P15. Red, P13; blue, P15.Scale bars: 20 pA (vertical red), 10 pA (vertical, blue), 50 mV (vertical, black) and 20 ms (horizontal). Fourpresynapticactionpotentials were evoked at 20 Hz. (**B**) Normalized peak amplitude of SST-IN **→**PC uIPSCs showed short-term depression, and nosignificant difference in PPR was found between P12–13 (red) and P14–15 (blue) mice. (**C**) Coefficient of variation (C.V.) in SST-IN **→** PC connections did not changed from P12–13 to P14–15. (**D**) The failure rate in SST-IN**→**PC connections did not change from P12–13to P14–15. (**E**) Representative uIPSC responses from a layer 2/3 SST-IN to a layer 2/3 PC evoked by a train of 3 presynaptic actionpotentials at 20 Hz under three different external Ca^2+^/Mg^2+^ concentrations. The postsynaptic cells were recorded with Cs-based and highCl-intracellular solution. Scale bars: 100 pA (vertical) and 20 ms (horizontal). (**F**) Parabola fit plot of the variance and mean of peakamplitude of SST-IN**→**PC uIPSCs in (**E**) under three different external Ca^2+^/Mg^2+^ concentrations. Red dots, 1 mM Ca^2+^/3 mM Mg^2+^;green dots, 2 mM Ca^2+^/2 mM Mg^2+^; blue dots, 3.7 mM Ca^2+^/0.3 mM Mg^2+^. (**G**) The number of release sites in SST-IN**→**PC connectionsdid not change from P12–13 to P14–15. (**H**) The quantal size in SST-IN**→**PC connections significantly decreased from P12–13 toP14–15. (**I**) Amplitude-scaled overlay of paired-pulse ratio (PPR) responses in FS-IN**→**PC connections at P13 and P15. Red, P13; blue, P15. Scale bars: 100 pA (vertical red and blue), 50 mV (vertical, black) and 20 ms (horizontal). (**J**) PPR in FS-IN**→**PC connections wassimilar between P12–13 (red) and P14–15 (blue) mice. (**K**) Coefficient of variation (C.V.) in FS-IN**→**PC connections was unchangedfrom P12–13 to P14–15. (**L**) The failure rate in FS-IN**→**PC connections did not change from P12–13 to P14–15. (**M**) RepresentativeuIPSC responses from a layer 2/3 FS-IN to a layer 2/3 PC evoked by a train of 3 presynaptic action potentials at 20 Hz in differentexternal Ca^2+^/Mg^2+^ concentrations. Scale bars: 200 pA (vertical) and 20 ms (horizontal). (**N**) Parabola fit plot of the variance and mean ofuIPSC amplitude in (**M**) at different external Ca^2+^/Mg^2+^ concentrations. (**O**) The number of release sites in FS-IN**→**PC connections didnot change from P12–13 to P14–15. (**P**) The quantal size in FS-IN**→**PC connections significantly increased from P12–13 to P14–15. Detailed statistical analysis, detailed data and exact sample numbers are presented in the ***Figure 7*-*source data 1.*** Error bars indicatemean ± SEM. *, *p* < 0.05; ***, *p* < 0.001; *n.s.* for *p* > 0.05.

Similarly, in FS-IN→PC synaptic transmission, there were no significant differences in PPR, C.V., and failure rate between P12–13 and P14–15 mice (***Figure 7I, 7J, 7K*** and ***7L***). Moreover, the average values of N at P12–13 and P14–15 were not significantly different (P12–13, 12.9 ± 3.1, n = 8; P14–15, 15.8 ± 1.9, n = 8; two-tailed unpaired *t* test, *p* = 0.439, ***Figure 7O***). Unlike SST-IN→PC connections, Q of FS-IN→PC connections at P14–15 was significantly larger than that at P12–13 (P12–13, 11.5 ± 1.9 pA, n = 8; P14–15, 27.5 ± 6.2 pA, n = 8; two-tailed unpaired *t* test, *p* = 0.026, ***Figure 7P***).

Together, these results suggest the postsynaptic quantal sizes in both of SST-IN→PC and FS-IN→PC synaptic transmission are altered during eye opening.

## Discussion

The plasticity of GABAergic circuitry in the visual critical period in the developing visual cortex has been extensively studied by manipulations that disrupt normal visual experience after eye opening (***Griffen and Maffei, 2014***; ***Hensch, 2005***; ***Lefort et al., 2013***; ***Maffei et al., 2004, 2010***). However, the changes and plasticity of GABAergic circuitry during eye opening, a natural event in the course of animal development, are still far from fully understood (***Gandhi et al., 2005***; ***Kuhlman et al., 2011***; ***Maffei et al., 2004***). In this study, we show that (1) eye opening weakens the synaptic transmission from SST-INs to PCs, but increases the synaptic transmission from FS-INs to PCs, (2) the inhibitory synaptic transmission from SST-INs to other types of interneurons remains unaltered during eye opening, (3) eye opening-induced alteration of the inhibitory synaptic transmission onto PCs is mediated by changes in postsynaptic quantal size.

We studied the formation of inhibitory synapses onto layer 2/3 PCs and focused on SST-INs and FS-INs. Although *SST*-*IRES*-*Cre* line has been widely used to study SST interneurons in cortical layer 2/3 both *in vivo* and *in vitro*, the neurons targeted by this line are heterogeneous (***Hu et al., 2013***; ***Jiang et al., 2015***). Indeed, we observed that ∼5.2% of SST-tdTomato neurons in cortical layer 2/3 of visual cortex express PV and ∼17.2% of SST-tdTomato neurons show the fast-spiking properties. In spite of this heterogeneity, after removing these fasting-spiking tdTomato^+^ neurons, we observed the vast majority of tdTomato^+^ cells in cortical layer 2/3 of the neocortex are the Martinotti cell subtype (95%, 19 out of 20). Taking advantage of *SST*-*tdTomato::Lhx6*-*EGFP* line, FS-INs were identified by EGFP^+^/tdTomato^-^ and fast-spiking properties in this study. Unlike PV-INs that innervate the cell body and proximal dendrites of the layer 2/3 PCs, SST-INs innervate the distal regions of PCs, including the apical dendrites ***(Chen et al., 2015***; ***Cristo et al., 2004***). Notably, the synaptic strength measured in our study represents the strength at the soma of the recorded neuron rather than at the contact sites. These somatic recordings undoubtedly underestimate the distal dendritic tonic currents due to attenuation within the dendrites and limited spatial reach of somatic voltage clamp (***Williams and Mitchell, 2008***). Given that there are no significant differences in both the total length and complexity of PC dendrites (***Figure 1-figure supplement 3***), and the rise time and half-width of uIPSCs between P12–13 and P14–15 mice (***Figure 1F*** and ***1G***), the relative change in the specific SST-IN→PC connection strength is unlikely due to space-clamp bias. Moreover, we observed the connection probability and strength of SST-IN→PC uIPSCs exhibit similar developmental properties when we used a cesium-based intracellular solution (improve space clamp) containing high concentration of Cl^-^ (increase the driving force of uIPSCs) to record the postsynaptic currents.

We systematically examined the development of synaptic connections from SST-INs onto PCs in layer 2/3 of the visual cortex. We observed that SST-IN→PC connections emerge at P7–8, and the probability increases from P7–8 to P9–11. After P11, we found a dense and stable innervation of SST-INs onto PCs, which reinforces their potentially crucial role in network activity. The increase in connection probability (from P7–8 to P9–11) is prior to the decline in peak amplitude of SST-IN→PC uIPSCs (from P12–13 to P14–15), suggesting that the morphological growth and synapse function are two independent processes. The average probability of SST-IN→PC connections observed (∼58% within 100 µm at P12–20) is lower than previously described with two-photon photostimulation in layer 2/3 of the frontal cortex (∼70% within 200 µm) (***Fino and Yuste, 2011***) and with paired recordings in layer 2/3 of the visual cortex (∼100% within 100 µm) (***Pfeffer et al., 2013***), and higher than reported with double or triple patch-clamp recordings in layer 2/3 of the visual cortex (∼20%) (***Thomson and Lamy, 2007***; ***Yoshimura and Callaway, 2005***), in layer 4 of the visual cortex (∼40% at P14–15 and ∼20% at P20–22) (***Miao et al., 2016***) and in layer 5 of the visual cortex (∼3%) (***Otsuka and Kawaguchi, 2009***); but agrees with paired recordings in layer 2/3 of the somatosensory cortex (∼63%) (***Xu et al., 2013***). The discrepancies may be due to regional differences, layer specificity or methodological differences.

In addition, we observed that the development of inhibitory synaptic transmission onto PCs displays several distinct features in layer 2/3 of mouse neocortex. First, the strength of SST-IN→PC synaptic connections is rapidly reduced by ∼65% from P12n 13 to P14–15. Although the developmental synaptic transmission from SST-INs onto PCs has not been quantified systematically, previous work reported that connection strength from low-threshold spiking interneurons (putative SST-INs) to excitatory spinal neurons increases from P12–13 to P14–15 in layer 4 of the rat somatosensory cortex (***Long et al., 2005***). In contrast, a recent study found that both the connection probability and strength of SST-IN→PC synaptic connections decrease substantially from P14–15 to P20–22 within layer 4 of the mouse visual cortex (***Miao et al., 2016***). The discrepancies may be due to layer differences (see below). Unlike SST-IN→PC pairs, our data showed that the strength of FS-IN→PC synaptic inputs in layer 2/3 of visual cortex is rapidly increased by ∼140% from P12–13 to P14–15. However, previous study reported that FS-IN→PC unitary conductance remains unaltered after P8 in layer 5/6 of mouse visual cortex (***Pangratz-Fuehrer and Hestrin, 2011***). Similarly, Yang *et al*. observed the strength of FS-IN→PC uIPSCs is not changed after P9 in layer 5/6 of mouse prefrontal cortex (***Yang et al., 2014***). These studies imply that the FS-IN→PC synaptic transmission does not change in the cortical layer 5/6 during eye opening. Second, rapid change of synaptic transmission from SST-INs and FS-INs onto PCs coincides with the onset of eye opening. It should be noted that, while the changes of inhibitory synaptic transmission coincide with natural eye opening, we cannot determine the exact temporal sequence of the two events due to the variability in the timing of eye opening (1–2 d) and synaptic responses. Nonetheless, with a controlled eye opening paradigm (***Lu and Constantine-Paton, 2004***; ***Yoshii et al., 2003***), it will be interesting to further explore the sequence of events within the first 24 hours after eye opening. Lastly, the weakening of SST-IN→PC synaptic transmission during eye opening exists not only in layer 2/3 of the visual cortex, but also in layer 2/3 of the prefrontal Cg1/2 area. Indeed, eye opening has been shown to affect hippocampal development, and early eye opening accelerated the maturation of synaptic strength (***Dumas, 2004***).

A major finding from our recordings in layer 2/3 of the visual cortex is that natural eye opening regulates synaptic transmission from SST-INs and FS-INs onto PCs. We draw this conclusion based on three lines of experimental evidence. We found that visual deprivation (dark rearing) at an early postnatal period can prevent the weakening of SST-IN→PC synaptic transmission. Furthermore, we demonstrated that eye opening deprivation (binocular lid suture) can efficiently prevent the changes in both of SST-IN→PC and FS-IN→PC synaptic transmission. More importantly, we controlled the timing of eye opening by artificially opening the lids from the binocular lid-sutured mice two days after natural eye opening (P16) and compared synaptic responses between siblings with and without eye opening. Our data showed that artificially opening the eyes sufficiently induces the decrease of SST-IN→PC synaptic transmission and the increase of FS-IN→PC synaptic transmission. These results strongly suggest that eye opening can specifically regulate the SST-IN→PC and FS-IN→PC synaptic transmission in layer 2/3 of the visual cortex. Of note, a recent study found that although synaptic strength of SST-IN→PC connection significantly decreases from P14–15 to P20–22 within layer 4 of the mouse visual cortex, dark rearing has no effect on it (***Miao et al., 2016***). Moreover, it has been reported that brief monocular visual deprivation during the time of eye opening selectively reduces the strength of synaptic transmission from PV-INs to PCs in layer 4 of the visual cortex (***Maffei et al., 2004***). These results suggest that visual deprivation-induced change in GABAergic circuits is layer- and cell type-specific. Indeed, accumulating evidence suggests that there are striking differences in morphology, intrinsic electrophysiological properties and synaptic connectivity between layer 2/3 and layer 4 SST-INs (***Ma et al., 2006***; ***Xu et al., 2013***).

Interestingly, we observed that the peak amplitude of uIPSCs from SST-INs onto FS-INs and Htr3a-INs does not change during eye opening. Moreover, eyelid suture doesn t change the strength of synaptic transmission from SST-INs to FS-INs. These results suggest that within layer 2/3 of the visual cortex, the inhibitory synapses made by SST-INs exhibit remarkable specificity in their strength with specific targets during eye opening. It should be noted that Htr3a-INs are heterogeneous, and encompass a diverse pupulation of interneurons including VIP interneuorns, NPY interneurons and other types of interneurons (***Chittajallu et al., 2013***; ***Lee et al., 2010***; ***Pfeffer et al., 2013***). It will be interesting to further investigate the development of synaptic transmission from SST-INs to subtypes of Htr3a-INs during eye opening.

The physiological roles for the differential alterations of inhibitory synaptic transmission onto PCs during eye opening remain to be determined. Accumulated evidence suggests that SST-INs that densely innervate nearby PC dendrites in mouse cortical layer 2/3 are responsible for controlling the efficacy and plasticity of synaptic inputs (***Chen et al., 2015***; ***Chiu et al., 2013***). Given that in early postnatal life, GABAergic transmission is excitatory to immature postsynaptic neurons (***Ben-Ari, 2002***; ***Owens et al., 1996***), early emergence of SST-IN→PC synaptic transmission may enhance the excitability of PCs, thereby promoting their maturation and synaptogenesis (***Oh et al., 2016***). However, around the time of eye opening, SST-INs would inhibit PCs. Decreased SST-IN→PC inhibition after eye opening could therefore enhance the effect of visual input onto excitatory neurons in the visual cortex by facilitating dendritic events in distal regions (***Figure 8***). Contrary to SST-INs, FS-INs control the spike output of PCs by inhibiting their perisomatic sites. Increased FS-IN→PC inhibition after eye opening is a homeostatic response to the reduction of SST-IN inhibition and the resulting increase in the excitability of PCs (***Bloodgood et al., 2013***; ***Chen et al., 2015***). We speculate such enhanced visual input to the visual cortex gated by SST-INs and the homeostatic rebalancing of inhibition regulated by FS-INs might be important for visual integration, an essential step in visual perception.

**FIGURE 8:**
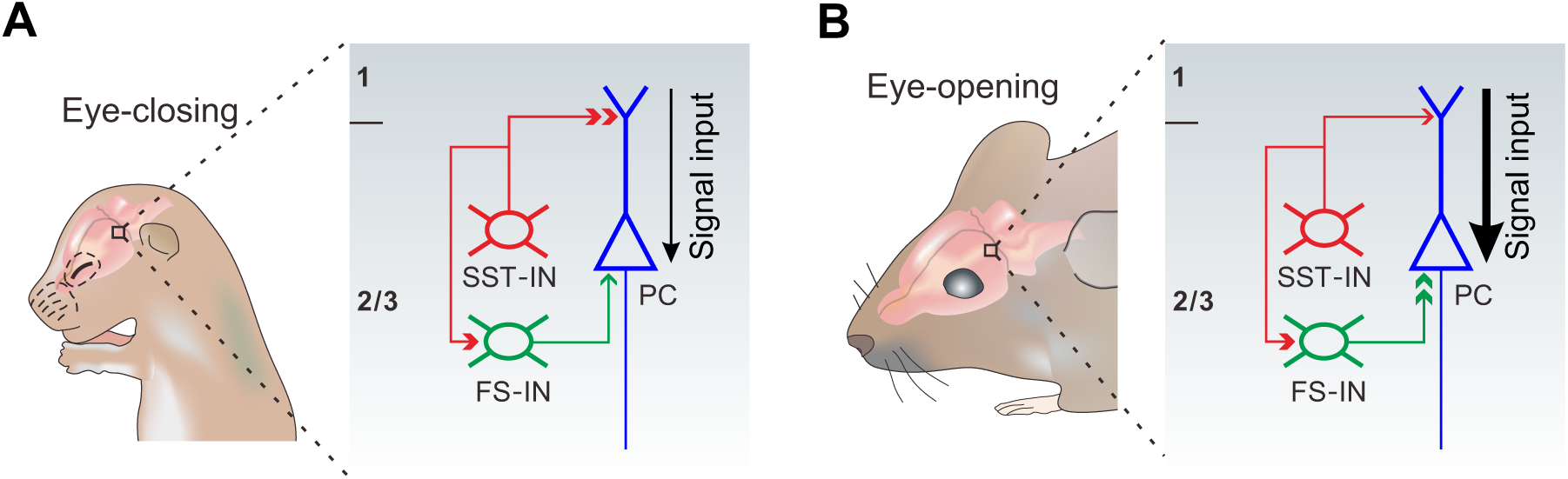
A model depicting how eye opening differentially regulates inhibitory synaptic transmission in developing layer 2/3 of the visual cortex. The development of inhibitory synaptic transmission before (A) and after eye opening (B). SST-INs primarily innervate distal dendrites of PCs, while FS-INs mainly target and inhibit somatic and perisomatic regions of PCs. Eye opening decreases the inhibition from SST-INs to PCs. Decreased SST-IN→PC inhibition after eye opening is predicted to enhance the effect of visual input onto excitatory neurons in the visual cortex by facilitating dendritic events. In contrast, synaptic inputs from FS-INs to PCs increase after eye opening. The increase of FS-IN→PC inhibition is speculated to involve in a homeostatic rebalancing of inhibition. The inhibition from SST-INs to FS-Ins remains unchanged during the time of eye opening.

## Materials and methods

### Animals

Four transgenic mouse lines, including *SST*-*IRES*-*Cre* (JAX no.13044), *Htr3a*-*EGFP* (MGI: 3846657), *Lhx6*-*EGFP* (MGI: 3839374) and tdTomato reporter *Ai14* (JAX no.007914) were used in this study. The *SST*-*IRES*-*Cre* mice were crossed to *Ai14* mice to generate *SST*-*tdTomato* alleles. The *Htr3a*-*EGFP* and *Lhx6*-*EGFP* mice were crossed to *SST*-*tdTomato* mice to generate *SST*-*tdTomato::Htr3a*-*EGFP* and *SST*-*tdTomato::Lhx6*-*EGFP* alleles. All pups were reared under a normal 12-hour light/dark cycle, unless otherwise stated. The day of parturition was defined as postnatal day 1 (P1). Pups were examined daily to monitor the postnatal day of eye opening. All experiments followed the guidelines for the care and use of laboratory animals at Fudan University.

### Dark rearing and eyelid suture

For dark rearing, pups were raised in dark cages after P3 until sacrificed for in vitro recordings. For eyelid suture, P8 mice were first carefully anesthetized with isoflurane and disinfected with ethanol. The binocular eyelids were sutured with small sterile ophthalmic needles. For artificial eye opening, sutured mice were anesthetized with isoflurane and the eyelids were carefully opened at P16. After eyelid suture or artificial eye opening, the eyelids were covered with tetracycline ointment, and the pups were kept on warm blankets until fully recovered. If the eyelids of sutured mice were unexpectedly open before recording, the pups were discarded and not included in the recording experiment.

### Brain slice preparation

P5–20 mice were anaesthetized with 1% isoflurane and 0.5–1.0 L/min oxygen. Brains from P5 to P8 mice were cut coronally at a thickness of 300 μm with a Compresstome VF-300 (Precisionary Instruments, USA) in a chilled solution containing (in mM) 120 choline chloride, 2.6 KCl, 26 NaHCO_3_, 1.25 NaH_2_PO_4_, 15 glucose, 1.3 ascorbic acid, 0.5 CaCl_2_ and 7 MgCl_2_ (pH 7.3–7.4, 300–305 mOsm), then incubated in artificial cerebrospinal fluid (ACSF) containing (mM) 126 NaCl, 3 KCl, 1.25 KH_2_PO_4_, 1.3 MgSO_4_, 3.2 CaCl_2_, 26 NaHCO_3_ and 10 glucose (pH 7.3–7.4, 300–305 mOsm), bubbled with 95% O_2_/5% CO_2_. P9 to P20 brain slices were prepared using a protective slicing and recovery method reported previously (***Zhao et al., 2011***). Briefly, anaesthetized mice were perfused intracardially with ice-cold oxygenated (95% O_2_, 5% CO_2_) NMDG-based cutting solution containing (in mM) 93 NMDG, 2.5 KCl, 1.2 NaH_2_PO_4_, 30 NaHCO_3_, 20 HEPES, 25 glucose, 5 sodium ascorbate, 2 thiourea, 3 sodium pyruvate, 10 MgSO_4_, 0.5 CaCl_2_ and 12 NAC (pH 7.3–7.4, 300–305 mOsm). Brains were carefully removed from the skull, cut coronally at a thickness of 300μm with a Compresstome VF-300 in chilled oxygenated (95% O_2_, 5% CO_2_) NMDG-based cutting solution. Slices were initially recovered in NMDG-based cutting solution at 32°C for 10 mins. Slices were then incubated in oxygenated (95% O_2_, 5% CO_2_) HEPES-modified solution containing (in mM) 94 NaCl, 2.5 KCl, 1.2 NaH_2_PO_4_, 30 NaHCO_3_, 20 HEPES, 25 glucose, 5 sodium ascorbate, 2 thiourea, 3 sodium pyruvate, 2 MgSO_4_, 2 CaCl_2_ and 6 NAC (pH 7.3–7.4, 300–305 mOsm) at room temperature for 40 mins. Finally, slices were incubated in oxygenated (95% O_2_, 5% CO_2_) normal ACSF at room temperature for at least 1h before recording.

### Electrophysiological recording and analysis

Slices were transferred to a recording chamber, which was constantly perfused with fresh normal ACSF at 32–34 °C, bubbled with 95% O_2_/5% CO_2_. Cells were visualized with water immersion objective (x20 and x60) and a BX51XI infrared-DIC microscope (Olympus, Japan) equipped with epifluorescence illumination. Glass recording electrodes (6-10 MΩ resistance) were filled with an intracellular solution consisting of (in mM) 93 K-gluconate, 16 KCl, 2 MgCl_2_, 0.2 EGTA, 10 HEPES, 2.5 MgATP, 0.5 Na_3_GTP, 10 Na-phosphocreatine, 0.4% neurobiotin (Invitrogen, USA) and 0.25% Alexa 568 or Alexa 488 (Invitrogen, USA) (adjusted to pH 7.25 and 295 mOsm). In some experiments, postsynaptic currents were recorded with Cs-based intracellular solution containing (in mM) 65 cesium methanesulfonate, 60 cesium chloride, 10 HEPES, 4 MgATP, 0.3 Na_3_GTP, 0.5 EGTA, 10 Na-phosphocreatine, 0.4% neurobiotin and 0.25% Alexa 488 (adjusted to pH 7.25 and 295 mOsm). Whole-cell recordings were obtained and analyzed using two Axon Multiclamp 700B amplifiers, Digidata 1440A (Molecular Devices, USA) and pCLAMP10 software (Molecular Devices, USA). Signals were sampled at 5 kHz with a 2 kHz low-pass filter. Images were captured with an ORCA-R2 digital CCD camera (Hamamatsu, Japan). To test synaptic connections, quadruple or triple whole-cell recordings were performed. To test unitary transmission, a brief suprathreshold current pulse (4–6 ms, 400–600 pA) was intracellularly injected to the presynaptic cell to elicit a presynaptic action potential, while the postsynaptic cells were held around −85 mV, and 20–30 repeated sweeps were recorded at time intervals of 10 or 20 s. After that, a train of 4 or 10 current pulses (20 Hz) were injected to the presynaptic cell to assay the efficacy of synaptic transmission. After recording, slices were fixed in 4% paraformaldehyde (PFA) in PBS overnight at 4°C.

Recordings with R_s_ > 30 MΩ were excluded from statistical analysis. R_s_ was compensated during recording. Electrophysiological data were analyzed off-line with Clampfit 10.6 (Molecular Devices). The rise time of uIPSCs was assayed from the 10% to 90% rising phase, and the half-width of uIPSCs was defined as the duration between the half amplitude. For the paired-pulse ratio calculation, the averaged peak amplitude of the first IPSC was defined as the basal level of synaptic strength. The C.V. was assayed from the amplitudes of each sweep. The failure rate was calculated as the percentage of sweeps without postsynaptic response. The fast-spiking physiological properties were identified as previously reported (***Hu et al., 2013***; ***Pangratz-Fuehrer and Hestrin, 2011***). For immature neurons (P5–1), fast-spiking properties were characterized by subthreshold oscillations ***(Pangratz-Fuehrer and Hestrin, 2011***).

The variance-mean analysis was performed as previously reported (***Mitra et al., 2012***; ***Scheuss and Neher, 2001***). Recordings were first carried out in ACSF containing 2 mM Ca^2+^/2 mM Mg^2+^, and then the chamber solution was changed to ACSF containing 3.7 mM Ca^2+^/0.3 mM Mg^2+^ and 1 mM Ca^2+^/3 mM Mg^2+^. Trains of 3 action potentials at 20 Hz were elicited and 30–40 repeated sweeps were recorded, with 10–20 s sweep-to-sweep interval. Recordings with stable baseline were used for analysis. The mean (*M*) and variance (*V*) of uIPSC amplitude were calculated for each pulse. The relationship between *M* and *V* was fitted to the parabola *V* = *QM* – *M*^2^/*N* (*Q*, quantal size; *N*, number of release sites). Quadratic regression was done with GraphPad Prism 5 software (GraphPad Software). Only recordings with *R*^2^ > 0.45 (*R*, regression index) were included for analysis.

### Immunohistochemistry and morphological reconstruction

Aneshetized P30 *SST*-*tdTomato::Htr3a*-*EGFP* and *SST*-*tdTomato::Lhx6*-*EGFP* mice were transcardially perfused with cold phosphate buffered saline (PBS), followed by 4% paraformaldehyde (PFA) in PBS. Brains were carefully removed from the skull, post-fixed overnight at 4°C. The brains were rinsed in PBS and sectioned into 60 μm thick coronal slices with a VT1000S vibratome (Leica, Germany). After that, free-floating slices were incubated with primary antibodies in blocking solution (1% bovine serum albumin, 0.5% Triton X-100 and 0.05% sodium azide in PBS) for 48 hours at 4°C. Slices were then washed with PBST (0.1% Triton X-100 in PBS) 5 times (10 min each), and incubated in blocking solution containing secondary antibodies for 24 hours at 4°C. The primary antibodies included mouse anti-RFP (1:500, #200301379, Rockland, USA), chicken anti-eGFP (1:1000, #1020, Aves, USA) and rabbit anti-parvalbumin (1:500, ab11427, Abcam, USA). The secondary antibodies were donkey anti-mouse (1:200, Alexa Fluor 555, A31570, Invitrogen, USA), donkey anti-chicken (1:200, DyLight 488, #703-546-155, Jackson ImmunoResearch, USA) and donkey anti-rabbit (1:200, Alexa Fluor 488, A21206, Life Technology, USA). For biocytin histochemistry, the fixed acute brain slices after electrophysiological recording were washed with PBS, incubated in blocking solution before incubation with Cy5-Streptavidin (1:500, #016-170-084, Jackson ImmunoResearch, USA) for 48 hours. Finally, sections were washed in PBS 5 times (10 min each), mounted and cover-slipped. Confocal images were taken using a FV1000 confocal microscope (Olympus, Japan) with 20x objective and 1 μm z-step size. Neurons were then reconstructed with Neurolucida Software (MicroBrightField, USA).

### Quantification and statistical analysis

Data were analyzed with SPSS 22 software (IBM) and GraphPad Prism 5 software (GraphPad Software). Statistical significance between groups was tested by two-tailed one-sample *t* test, two-tailed unpaired *t* test, paired *t* test, Mann-Whitney *U* test, Fisher s exact test, χ^2^ test, one-way ANOVA and two-way ANOVA. All the detailed test methods, the number of experiments and the *p* value are listed in the ***source data***. All data are presented as mean ± SEM, and difference was recognized as significant when *p* < 0.05.

## Acknowledgements

We thank Dr. Zhengang Yang for providing *Htr3a*-*EGFP* and *Lhx6*-*EGFP* mice. We thank Dr. Song-Hai Shi (MSKCC) for comments on the manuscript, and members of Y.-C.Y. laboratory for their discussions. This work was supported by grants from the National Key Research and Development Program of China (2016YFA0100802) and the Natural Science Foundation of China (31200816) to Y.F.; grants from the Natural Science Foundation of China (31271157, 31471036, 31629004, 31421091, 91332110), the Ministry of Science and Technology of China (2014CB942800 and 2012CB966303), the Foundation of the Ministry of Education of China (20130071110065), the Key Project of Shanghai Science and Technology Commission (No.15JC1400102) and Shanghai Municipal Science and Technology Committee of Shanghai outstanding academic leaders plan (15XD1500700) to Y.-C.Y.

## Author Contributions

W.G. and Y.-C.Y. conceived the project and designed the experiments. Y.F. and Y.-C.Y. supervised the project. J.-W.C. performed the morphological reconstructions and part of the electrophysiological recordings. Z.-H.Z performed part of the morphological reconstructions. W.G., J.-W.C. and Y.-C.Y. interpreted the data. W.G., Y.F. and Y.-C.Y. wrote the manuscript with comments from the rest of the authors.

---

**Figure 1-figure supplement 1:**
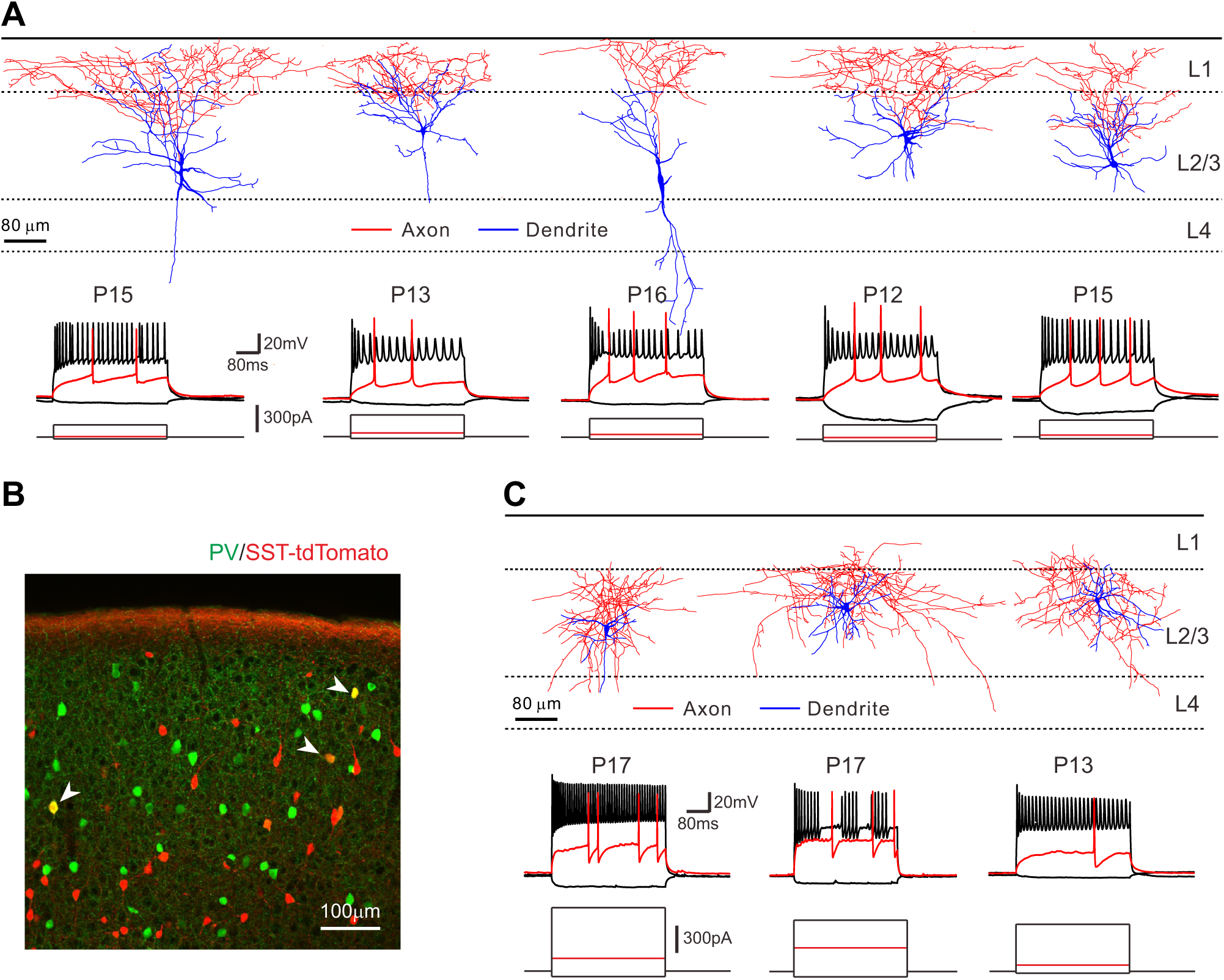
Morphological and electrophysiological properties of tdTomato *SST*-*tdTomato* mice. **(A)** Top panel, the morphologies of reconstructed non-FS tdTomato+ neurons in layer 2/3 of visual cortex. Bottom panel, corresponding traces of voltage responses to 500 ms current pulse step injections recorded in the current-clamp mode. (B) Some tdtomato+ cells in *SST*-*tdTomato* line expressed PV. (C) Top panel, the morphologies of reconstructed fast-spiking tdTomato+ cells. These cells are basket-like interneurons, with dense axonal arborlization in layer 2/3. Bottom panel, corresponding voltage membrane responses of cells to current injections.

**Figure 1-figure supplement 2:**
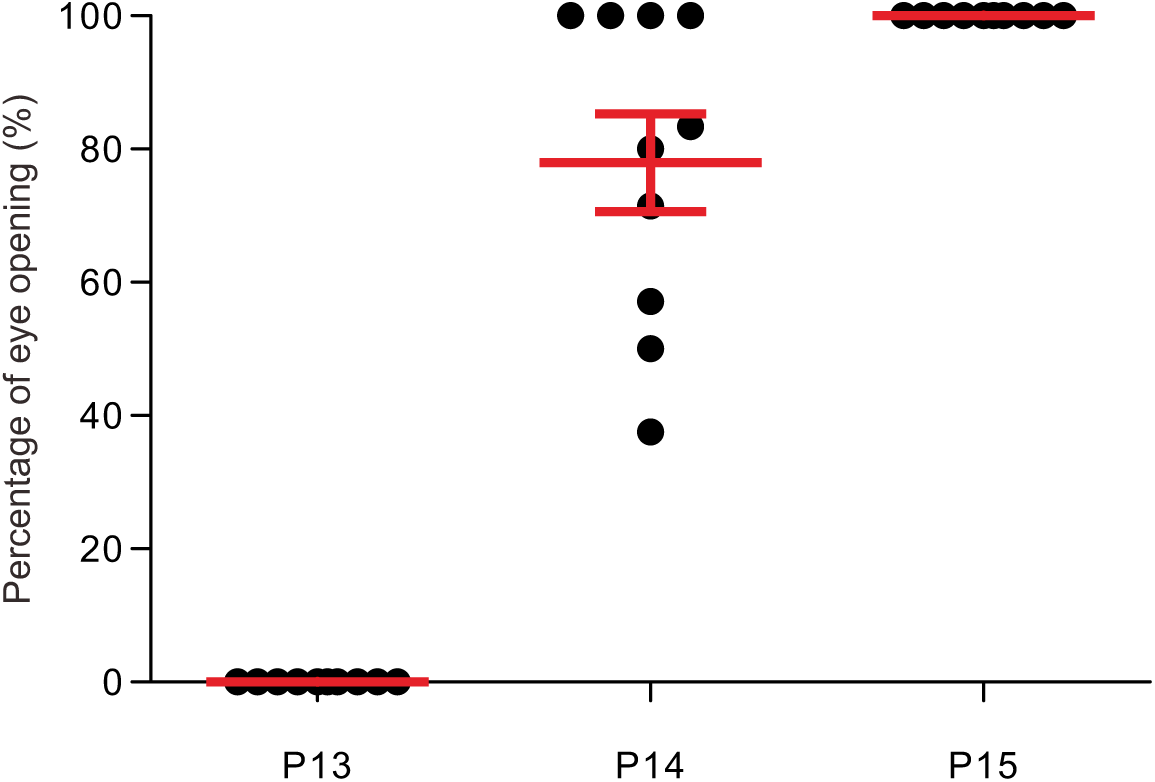
The timing of eye opening in mice. Quantification of mice undergoing eye opening in each litter (n = 10) from P13 to P15. There were mice with opened eyes at P13. At P14 77.9% ± 7.4% of mice had opened eyes and at P15 100% of mice had opened eyes.

**Figure 1-figure supplement 3:**
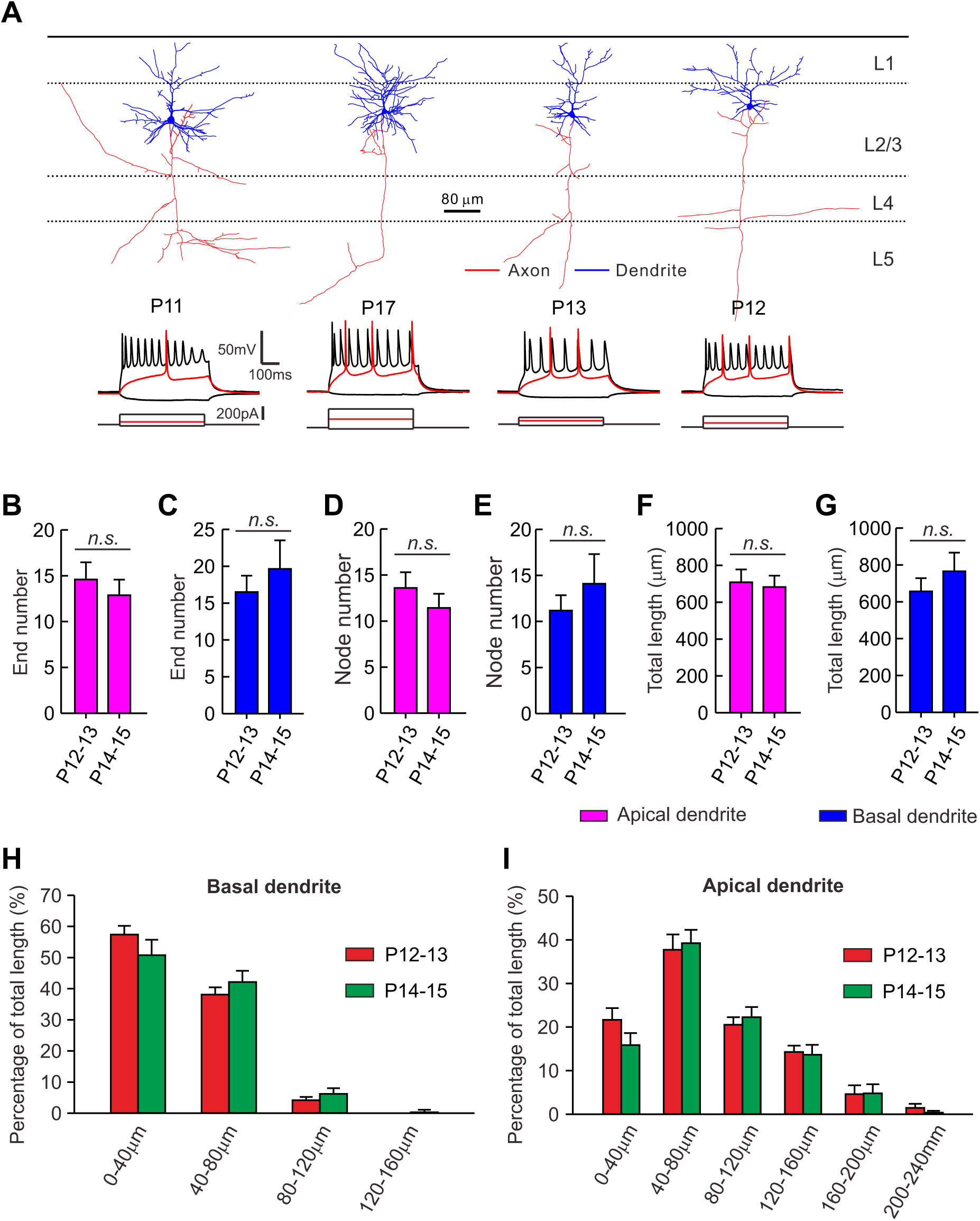
Dendritic morphology of PC_s_ does not change significantly before and after eye opening. **(A)** Morphological identity of recorded PCs. Top panel, the morphologies of reconstructed PCs. Bottom panel, corresponding membrane responses of PCs to current injections**. (B-G)**Quantification of the end number of apical dendrites **(B),** the end number of basal dendrites **(C)**, the node number of apical dendrites **(D),** the node number of basal dendrites **(E)**, the total length of apical dendrites **(F)** and the total length of basal dendrites **(G)** between P12–13 (before eye opening) and P14–15 (after eye opening). **(H-I)** Sholl analysis of dendritic length per 40mradialunit distance from soma. Detailed statistical analysis, detailed data and exact sample numbers are presented in the ***Figure 1*-*source data 1. n.s.*** for *p* > 0.05

**Figure 1-figure supplement 4:**
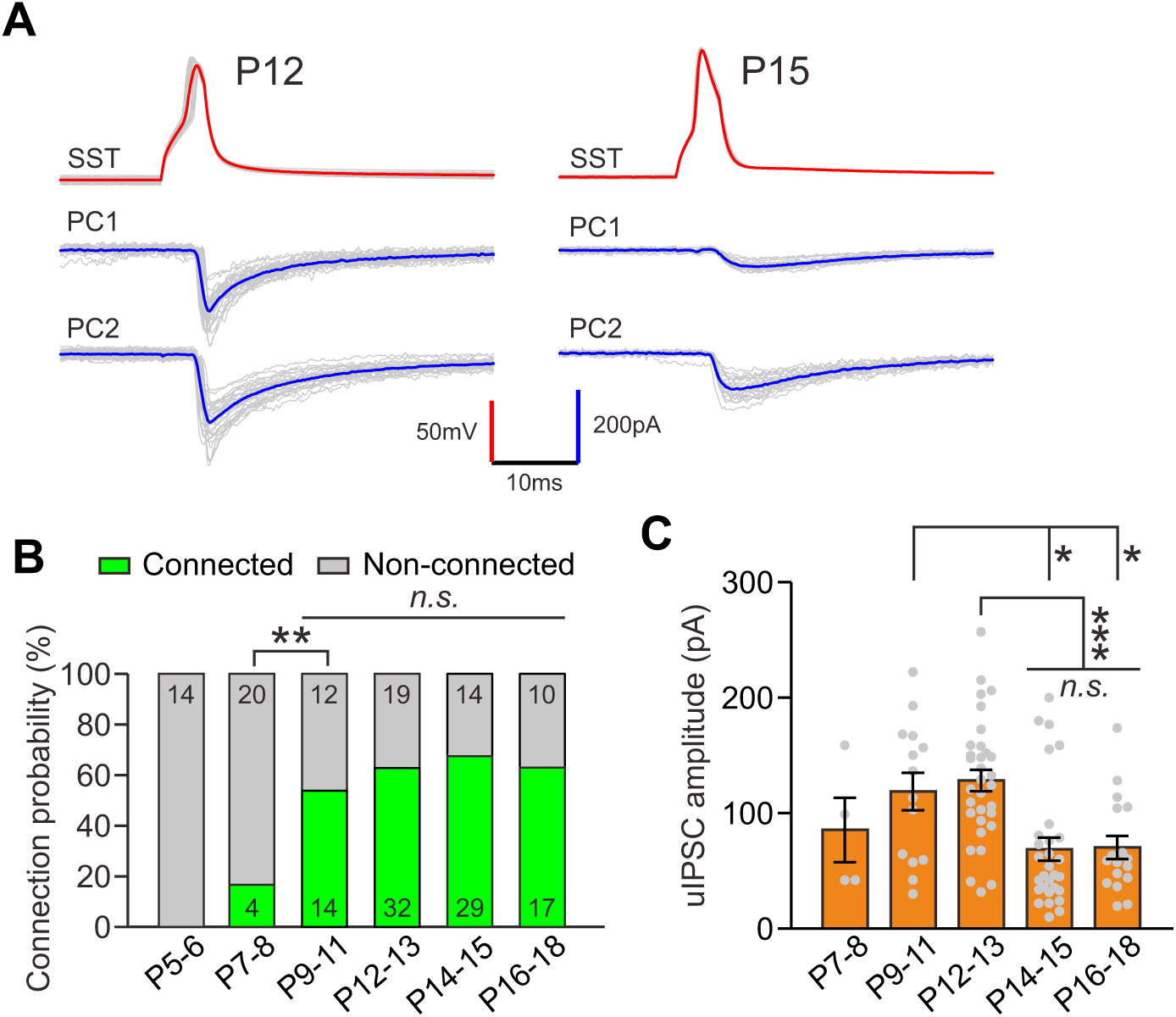
Synaptic responses from SST-INs to PCs recorded with cesium-based intracellular solution. (**A**)Representative traces showing synaptic transmission from SST-INs to PCs recorded at P12 andP15, respectively. Postsynaptic responses were recorded with cesium-based intracellular solutioncontaining 60 mM Cl-. **(B)** Summary of connection probability. **(C)** Quantification of peakamplitude of SST-IN→PC uIPSCs. Detailed statistical analysis, detailed data and exact sample numbers are presented in the ***Figure 1*-*source data 1***. *, *p* < 0.05; **, *p* < 0.01; ***, *p* < 0.001; *n.s.* for *p* > 0.05.

**Figure 3-figure supplement 1:**
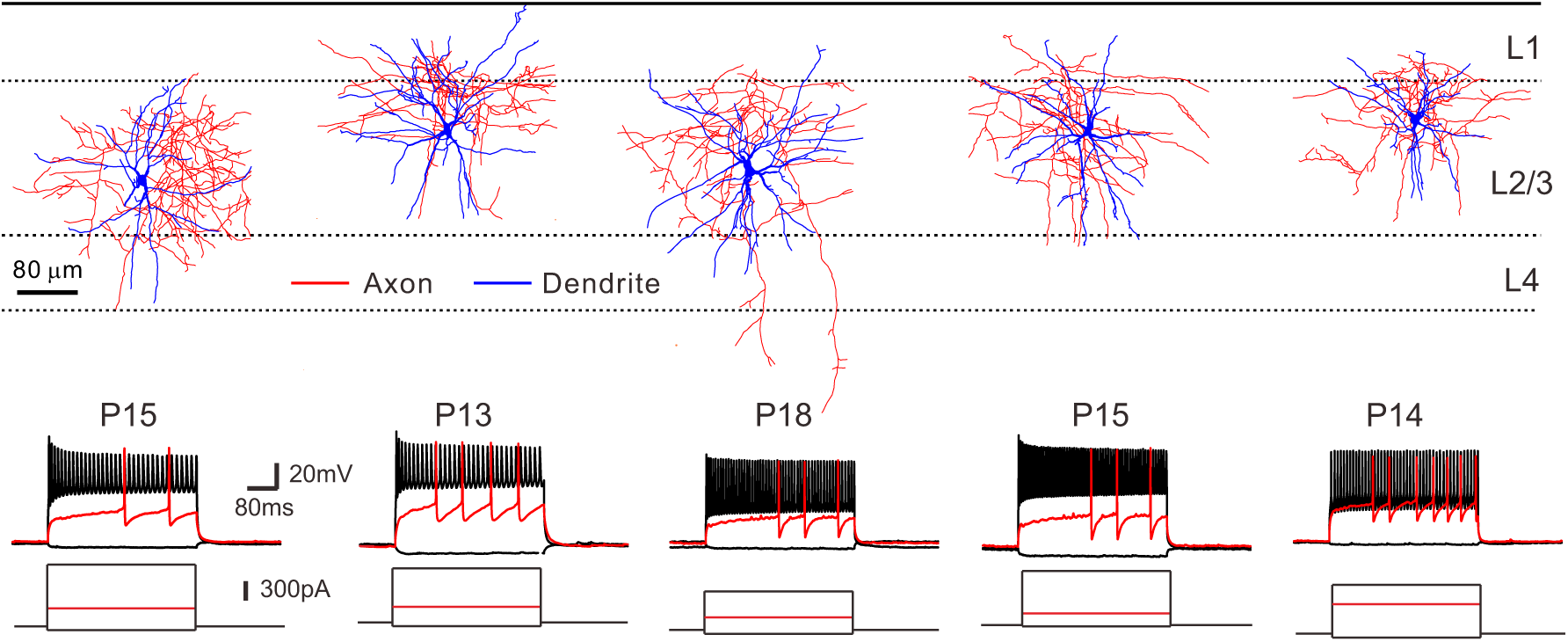
Morphologies of reconstructed FS-INs and corresponding evoked membrane responses. Top panel, the morphologic reconstructions of FS-INs. Bottom panel, the membrane responses ofFS-INs to current injections.

**Figure 6-figure supplement 1:**
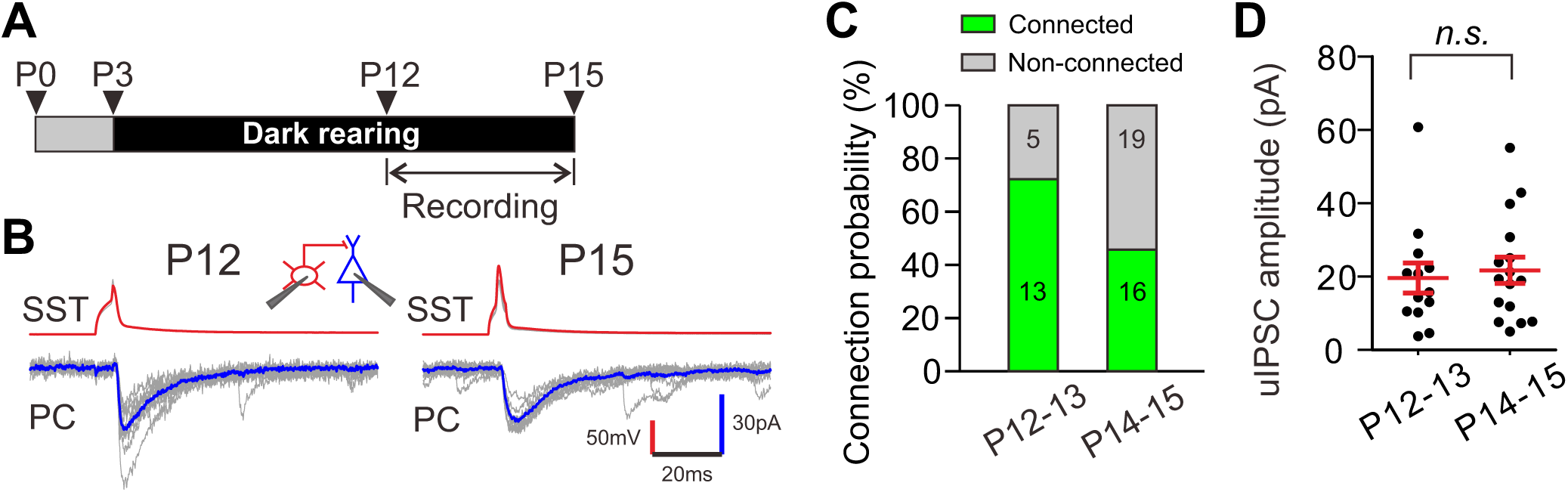
Dark rearing prevents the developmental decrease of synaptic transmission from SST-INs to PCs. (**A**) Schematic time schedule of dark rearing and electrophysiological recording. (**B**) Representative a layer 2/3 SST-IN to a layer 2/3 PC at P12 and P15 in dark-reared mice. Insert, schematic of paired recording of a SST-IN (red) and a PC (blue). (**C**) Occurrence of connection from layer 2/3 SST-INs to PCs in dark-reared mice was not significantly different between P12–13 and P14–15. (**D**) Peak amplitude of uIPSCs didn’t change significantly fromP12–13 to P14–15 in dark-reared mice. Detailed statistical analysis, detailed data and exact sample numbers are presented in the ***Figure 6*-*source data 1***. *n.s.* for *p* > 0.05.

**Figure 6-figure supplement 2:**
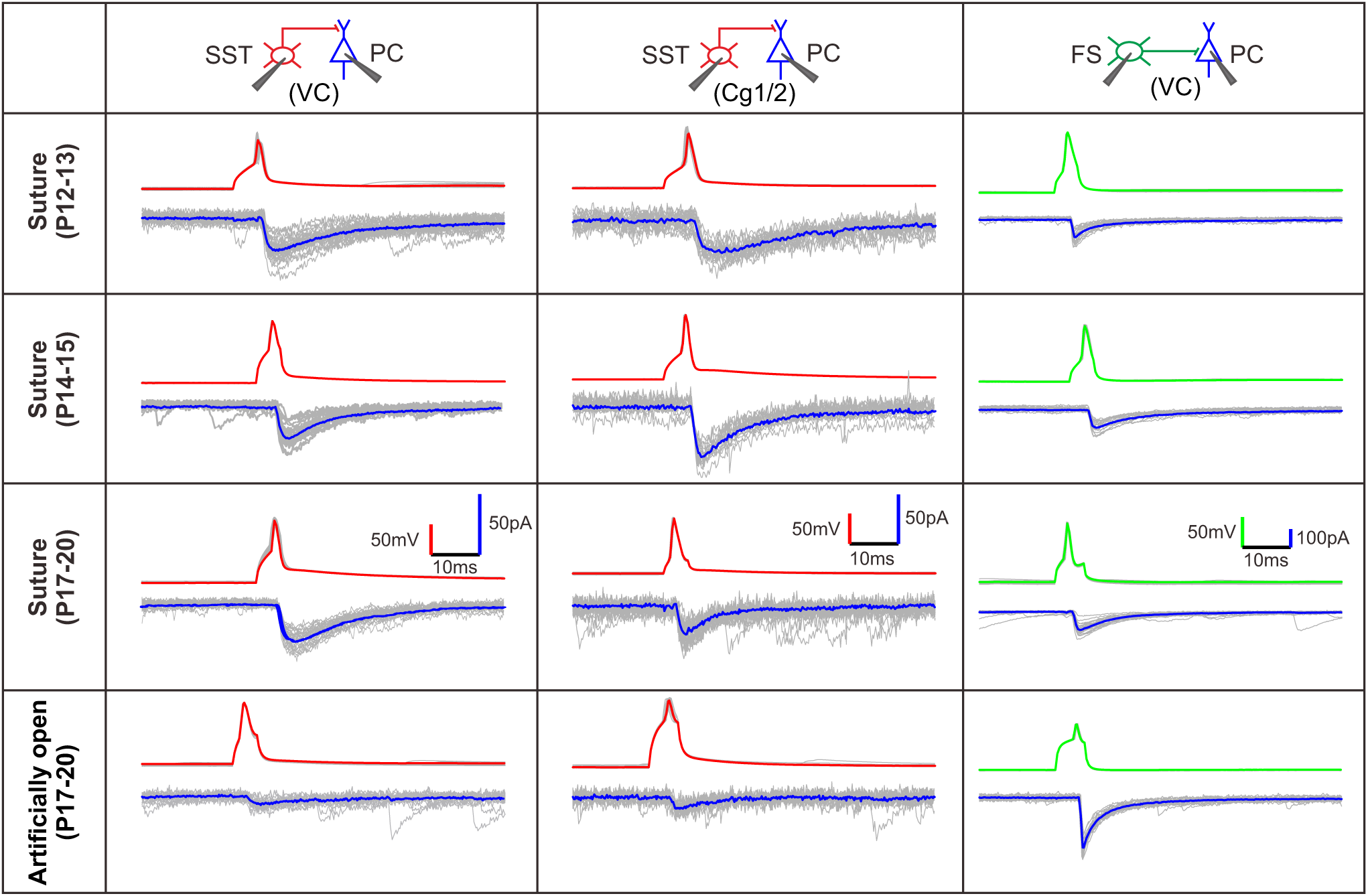
Representative traces of synaptic transmission (related to Figure 6). Top panels indicate the direction of synaptic transmission.

**Figure 6-figure supplement 3:**
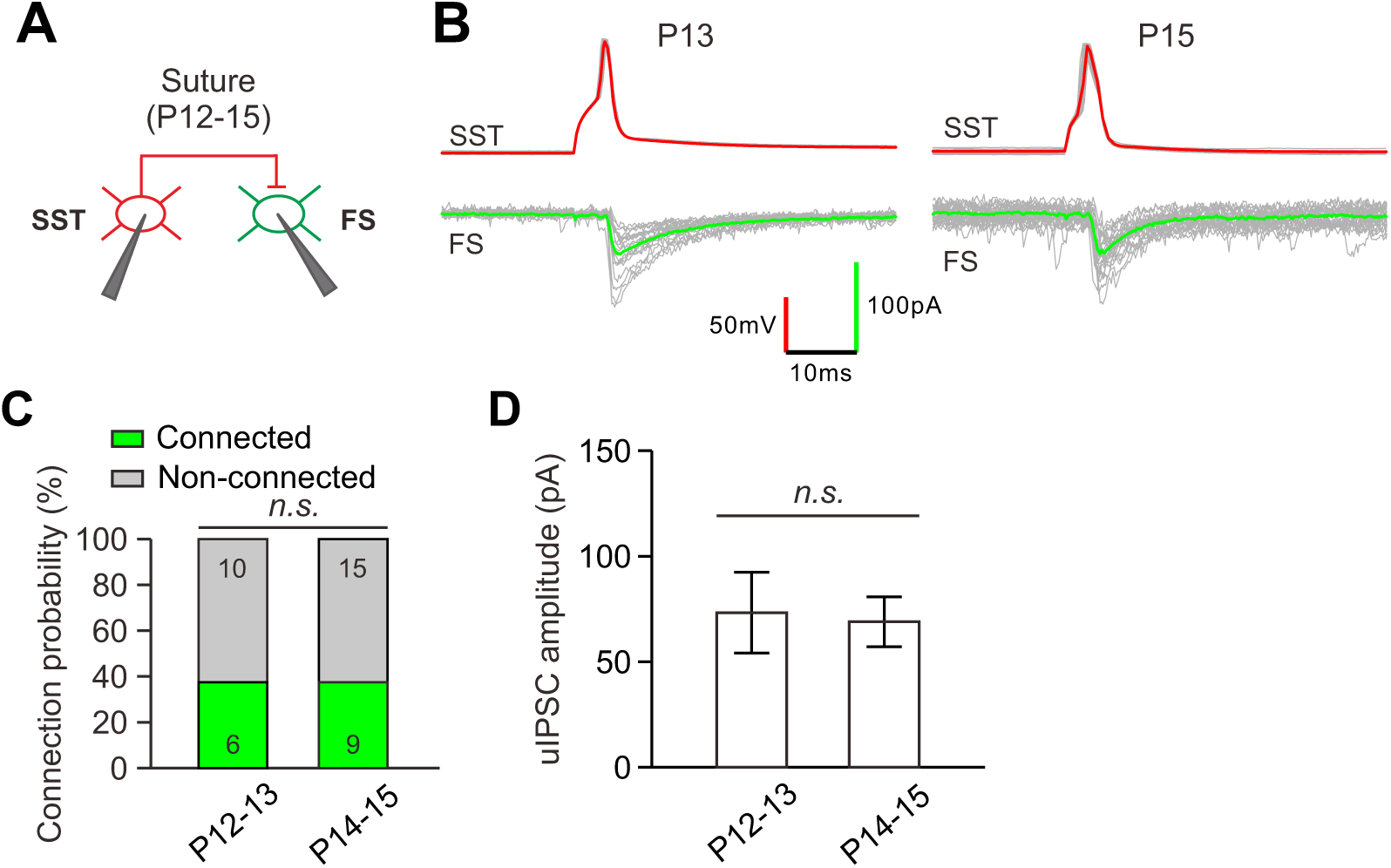
The strength of SST-IN →FS-IN synaptic transmission does not change in sutured mice during the time of eye opening. **(A)** Schema of a paired recording from a layer 2/3 SST-IN and a layer 2/3 FS-IN in sutured mice atP12–15**. (B)** Representative traces of synaptic transmission from a layer 2/3 SST-IN to a layer 2/3FS-IN at P13 and P15 in sutured mice. **(C)** Quantification of the connection probability. **(D)** Quantification of uIPSC peak amplitude. Detailed statistical analysis, detailed data and exact sample numbers are presented in the ***Figure 6*-*source data 1.*** *n.s.* for *p* > 0.05.

